# Bi-allelic mutations in *KCTD11* cause a new form of autosomal recessive intermediate Charcot-Marie-Tooth disease

**DOI:** 10.1101/2025.06.30.661538

**Authors:** Jihane Gadacha, Zahraa Haidar, Nathalie Roeckel-Trévisiol, Amandine Pauset, Christel Castro, Coralie Provost, Zeinab Hamzé, Camille Humbert, Karine Bertaux, Nicolas Lenfant, Marion Masingue, Alix de Becdelièvre, Marina Konyukh, Nathalie Bonello, Anne-Sophie Lia, Emilien Delmont, Alessandro Bertini, Ilaria Quartesan, Stefano Facchini, Andrea Cortese, Mary M Reilly, Henry Houlden, Davide Pareyson, Chiara Pisciotta, Shahram Attarian, Andoni Urtizberea, André Mégarbané, Rosette Jabbour, Nathalie Bernard-Marissal, Valérie Delague

**Author notes:** Correspondence to: Valérie DELAGUE, Marseille Medical Genetics, U 1251, Aix Marseille Université, Faculté de Médecine de la Timone, 27 bd Jean Moulin, 13385 Marseille cedex 05, France.

## Abstract

Charcot-Marie-Tooth disease (CMT) is the most common inherited neuromuscular disorder, characterized by progressive, length-dependent degeneration of peripheral nerves, resulting in distal muscle atrophy and weakness, foot and hand deformities, and sensory deficits. The disease is clinically and genetically heterogeneous, with over 125 disease-causing genes identified to date. Here, genetic studies in ten patients from 5 unrelated families of diverse ethnic background, led to the identification of KCTD11 as a novel CMT gene, responsible for a new autosomal recessive intermediate CMT subtype, RI-CMTE. The variants identified are loss of function.

*KCTD11* encodes KCTD11/REN, a protein of yet unknown function in the Peripheral Nervous System, known to regulate HDAC1, β-catenin, and mTORC1, key regulators of myelination and neuronal differentiation in the PNS.

To explore KCTD11’s role in the PNS, we used a constitutive Kctd11-/- mouse model and the derived in vitro myelin model of sensory neuron and Schwann cell co-culture (DRGN/SC), to mimic the loss-of-function induced by patient mutations. We first demonstrate that the loss of KCTD11 is due to enhanced degradation of the mutated protein via autophagy. Both in vitro and in vivo, we demonstrate abnormal myelination in vivo and altered myelination dynamics in vitro. These defects were associated with dysregulation of the expression of key transcription factors in Schwann cells, such as Egr2 and Sox10, along with other myelin-related genes, as revealed by mRNA-sequencing data. Regarding pathophysiological mechanisms, we identified dysregulation of HDAC1 expression, as well as alterations in the Wnt/β-catenin, Sonic Hedgehog and Hippo/YAP signaling pathways. The deregulation of these pathways seem to converge to altered autophagy and altered balance between proliferation, differentiation and apoptosis, at least in Schwann cells. These mechanisms remain to be explored in axons from PNS neurons.

Altogether, our results identify KCTD11 as a novel gene defective in autosomal recessive intermediate RI-CMTE and highlight the key role of KCTD11 in maintaining myelin homeostasis through regulation of HDAC1 and phosphorylated β-catenin levels, thereby preventing late-onset myelin abnormalities and degradation.

## Introduction

Charcot-Marie-Tooth (CMT), also known as Hereditary Motor and Sensory Neuropathy (HMSN), is one of the most common group of inherited neuromuscular disorders, with an estimated overall prevalence of 1 in 2,500 individuals (Skre 1974), which however can vary considerably between regions and populations, ranging from 1:1,250 to 1:10,000 (Barreto, Oliveira et al. 2016). This length-dependent peripheral neuropathy is characterized, clinically, by progressive muscular weakness starting at the distal extremities, foot deformities, loss of deep tendon reflexes, and mild to moderate distal sensory loss (Harding and Thomas 1980). Historically, two main subtypes have been defined based on histological and electrophysiological criteria: the demyelinating forms (CMT1 and CMT4), resulting from primary damage of myelinating Schwann cells, and characterized by slowed Nerve-Conduction Velocities (NCV, ≤38m/s in the upper limbs); and the axonal forms (CMT2), with primary degeneration of axons from motor and sensory neurons, and characterized by normal NCVs, but reduced amplitudes of action potentials (Pareyson and Marchesi 2009, Pisciotta and Shy 2018, Pisciotta and Shy 2023).

Finally, the intermediate form (I-CMT) refer to subtypes characterized by intermediate NCV ranging between 25 and 45 m/s at the median nerve, and combination of features observed in both CMT1 and CMT2 (Nicholson and Myers 2006) (Berciano, Garcia et al. 2017). In families affected with I-CMT, some affected individuals may have slowed motor conduction velocities in CMT1 range, while other have NCVs in the CMT2 range (Nicholson and Myers 2006, Berciano, Garcia et al. 2017).

CMT is also characterized by extensive genetic heterogeneity, with all modes of inheritance described and over 100 genes identified to date (Laura, Pipis et al. 2019, Pipis, Rossor et al. 2019). Although mutations in fivegenes (*PMP22*, *GJB1*, *MPZ*, *MFN2*, and *SORD*) account for more than 90% of established molecular diagnosis (at least in populations from Northern Europe and North America) (Pipis, Rossor et al. 2019, Pisciotta and Shy 2023), a notable proportion of patients with a clinical diagnosis of CMT remain genetically undiagnosed (Pipis, Rossor et al. 2019, De Grado, Serio et al. 2025). These statistics; however, vary widely among populations (Higuchi and Takashima 2023). Despite the high number of known genes, many patients remain without genetic diagnosis, therefore there are still many genes to be discovered.

The KCTD family of human proteins includes 21 members, of largely unknown function, containing a potassium channel tetramerization domain (KCTD). Several have been implicated in various diseases, including neurocognitive disorders (KCTD3; (Alazami, Patel et al. 2015)), neurodevelopmental diseases (KCTD7; (Kousi, Anttila et al. 2012, Krabichler, Rostasy et al. 2012, Farhan, Murphy et al. 2014, Metz, Teng et al. 2018)), bipolar disorder (KCTD12; (Schwenk, Metz et al. 2010)), autism and schizophrenia (KCTD13; (Golzio, Willer et al. 2012, Escamilla, Filonova et al. 2017)), movement disorders (KCTD17; (Mencacci, Rubio-Agusti et al. 2015)), and obesity (KCTD15; (Willer, Speliotes et al. 2009, Baranski, Kraja et al. 2018)). Notably, their involvement in these diseases is largely attributed to their role role in the ubiquitin-proteasome system in mammals by conferring substrate specificity during the ubiquitination process, crucial for maintaining protein homeostasis in all cell type (Wang, Argiles-Castillo et al. 2020). Among these proteins, KCTD11 is the best characterized member. It is expressed in a broad range of tissues, with particularly strong expression in peripheral nerve tissues (as evidenced by GTEx gene expression data). This monoexonic gene gives rise to two isoforms: a long 271 amino acid (aa) long isoform, lKCTD11 (NM_001363642), and a short isoform, sKCTD11 (NM_001002914;) of 232 aa (Correale, Pirone et al. 2011). *KCTD11*, also known as *REN*, was first described by (Gallo, Zazzeroni et al. 2002), who identified the gene as promoting neural cell differentiation during the early stages of Central (CNS) and Peripheral Nervous System (PNS) development by modulating the proliferation/differentiation balance and promoting axonal growth. Remarkably, this gene was later characterized as a tumor suppressor in medulloblastoma cancer due to its role in downregulating the Sonic Hedgehog pathway (Di Marcotullio, Ferretti et al. 2004, Argenti, Gallo et al. 2005).. Notably, KCTD11 described functions are explained by its intrinsic properties in regulating protein degradation. In fact, despite its name, it acts, through two distinct binding domains, as an adaptor protein enabling the ubiquitination of substrate proteins. KCTD11 can form homodimeric or heterodimeric complexes with other members of the KCTD protein family (De Smaele, Di Marcotullio et al. 2011). Its N-terminal BTB/POZ domain (bric-à-brac, tram-track, broad complex/poxvirus zinc finger) binds to Cullin3 (CUL3), the core component of the essential E3 ubiquitin ligase complex, while its C-terminal domain recruits target proteins that are subsequently ubiquitinated and degraded by the proteasome. Through this mechanism, KCTD11 regulates proteins and signaling pathways involved in a wide range of biological functions. Namely, KCTD11 regulates the protein levels of the Histone Deacetylase 1 (HDAC1; <u>(Canettieri, Di Marcotullio et al. 2010)</u>, which leads to/enable the downregulation of the Sonic Hedgehog pathway, as well as the levels of β-catenin from the Wnt/β-catenin pathway (Yang, Han et al. 2021). Moreover, the protein appears to downregulate the Mechanistic Target of Rapamycin Complex 1 (mTORC1) activity via mechanisms that remain to be further elucidated (Chen, Wang et al. 2018). Interestingly, mTORC1, HDAC1 and β-catenin are key regulators of signaling pathways involved in the development and function of the PNS, particularly in Schwann cells (Jacob, Christen et al. 2011, Makoukji, Shackleford et al. 2011, Tawk, Makoukji et al. 2011, Jacob, Lotscher et al. 2014), In particular, the binding of KCTD11 to β-catenin inhibits the Wnt pathway by interfering with the nuclear translocation β-catenin to nucleus and further inhibits the Hippo pathway, through altered translocation of YAP to the nucleus(Yang, Han et al. 2021).

We report here ten patients from five unrelated families affected with a new autosomal recessive intermediate Charcot-Marie-Tooth subtype: RI-CMTE. The patients present with distal amyotrophy and foot deformities, marked by a middle/late onset and slow progression. The NCVs at the median nerve are intermediate, ranging between 25 and 45 m/s, and are associated with reduced amplitudes in the lower limbs in all patients, as expected for intermediate CMT. Genetic studies revealed bi-allelic mutations in the *KCTD11* gene, which we propose as a novel defective in this RI-CMTE disease. This study aims to identify and characterize *KCTD11* as a novel causative gene underlying Charcot-Marie-Tooth disease. Functional analyses revealed that these variants result in loss-of-function at the protein level. Given the limited knowledge regarding the physiological role of KCTD11 in the PNS, we investigated its function using *in vitro* and i*n vivo* knockout models. Our findings demonstrate that KCTD11 ablation impairs myelination both *in vitro*, in dorsal root ganglion neuron/Schwann cell (DRGN/SC) co-cultures, and *in vivo*, in mouse peripheral nerves. Our results identify KCTD11 as a key regulator of peripheral nerve physiology, and reveal its deficiency as a novel genetic cause underlying I-CMT disease.

## Materials and methods

### Genetic Analysis

#### Samples

After informed consent was obtained from all individuals included in this study, EDTA blood samples were collected, and genomic DNA was extracted from lymphocytes with the use of standard methods. All protocols performed in this study complied with the ethics guidelines of the institutions involved.

#### Genome wide screen

A genome-wide screen has been performed at the Centre National de Genotypage (CNG, Evry, France), using 400 polymorphic microsatellite Single Tandem Repeat (STR) markers with an average intermarker distance of 10 cM on the following 25 individuals: F1-VI.3, F1-VI.4, F1-VII.1.2.3.5.6.7.8.10.11.12.14.16, F1-VIII.3.9.10.11.12.17.18.19.20 and F1-IX.1 and 2 (**Figure 1A**). Considering autosomal recessive transmission and the presence of inbred marriages in the family, the analysis was performed using homozygosity mapping (Lander and Botstein 1987), as previously described (Delague, Bareil et al. 2000).

**Figure 1.**
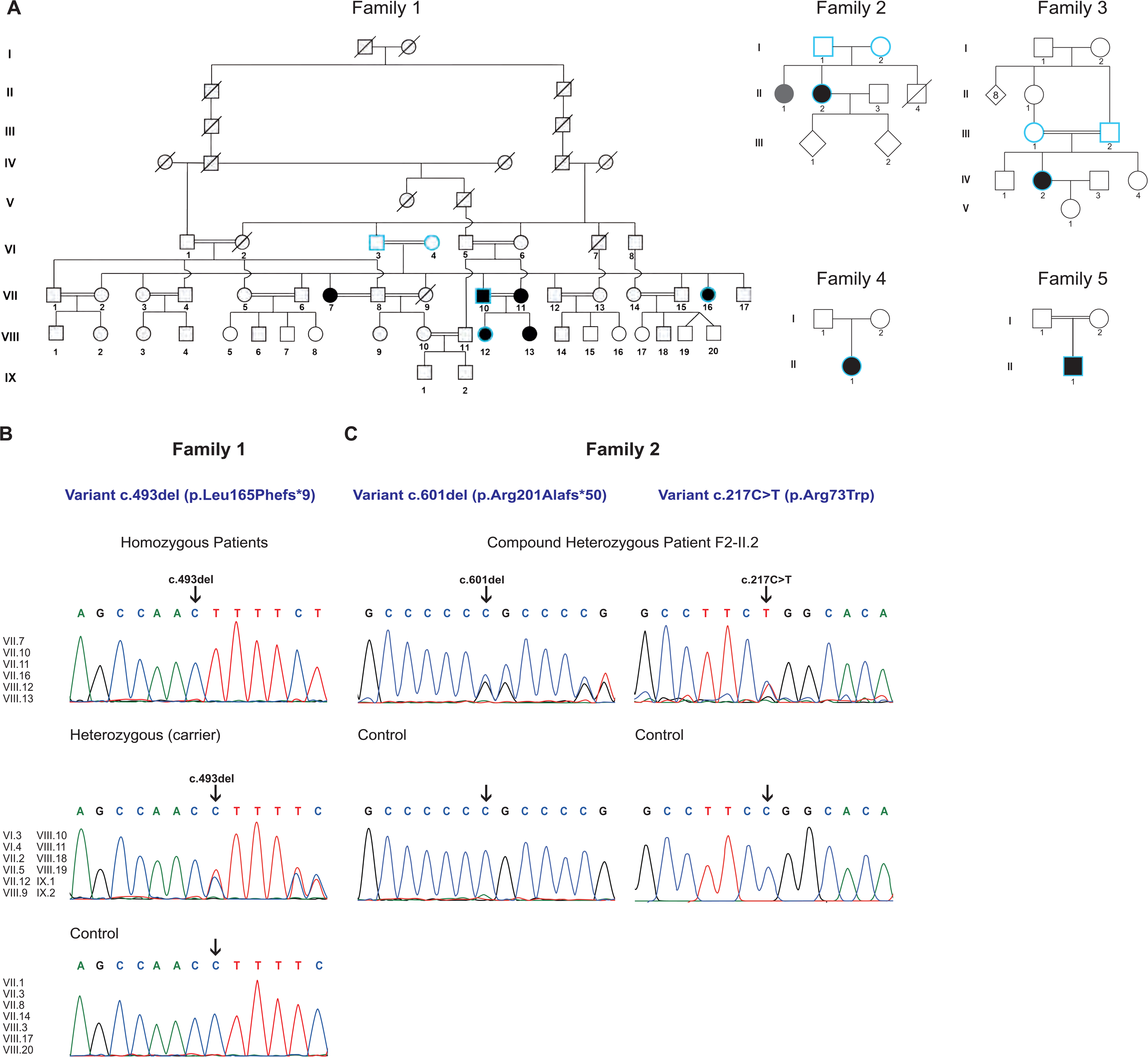
Genealogical and molecular characteristics of patients with mutations in *KCTD11*. (**A**) Pedigree of the five families affected with autosomal recessive intermediate CMT. Affected individuals are shaded in black. Individuals for whom the genome has been sequenced are highlighted in blue. The individual in gray has an unknown disease status. (**B-E**) Chromatograms of patients, unaffected parents, and controls showing the variants identified in *KCTD11* (NM_001363642). (**B**) Family 1: patients F1-VII.7, -VII.10, -VII.11, -VII.16, -VIII.12 and VIII.13 are homozygous for the c.493del (p.Leu165Phefs*9) variant. (**C**) Family 2: patient F2-II.2 is compound heterozygous for the following variants : c.601del (p.Arg201Alafs*50) and c.217C>T p.Arg73Trp). (**D**) Family 3: patient F3-IV.2 is homozygous for the c.310C>T (p.Gln104*) variant. . (**E**) Family 4: patient F4-II.1 is homozygous for the c.680G>C (p.Arg227Pro) missense variant. (**F**) Family 5: patient F5-II.1 is homozygous for the c.(269-282delinsC) (p.Tyr90Serfs*40) frameshift variant

#### Whole Genome Sequencing and analysis

##### Family 1

Whole genome sequencing was carried out on 5 individuals from Family 1 (F1-VII.10, F1-VII.16 and F1-VIII.12 and the unaffected parents F1-VI.3 and F1-VI.4). Library preparation, capture and sequencing were performed by the French National Genotyping Center (CNG, Evry, France), using a paired-end 100-bp read sequencing protocol. The quality of the reads was evaluated using fastQC 0.11.5. Raw data were mapped to the built of the human genome (hg19) by using BWA-MEM v0.7.12 (Li and Durbin 2009, Li 2013). Variant calling was subsequently performed using GATK v4.3.0.0 (McKenna, Hanna et al. 2010) and annotation was done with ANNOVAR (Wang, Li et al. 2010). All subsequent steps were performed using our *in-house* software for variant annotation and segregation VarAFT (Desvignes, Bartoli et al. 2018). WGS data from the five sequenced individuals were analyzed simultaneously and segregated using this tool. Considering autosomal recessive transmission and the high level of consanguinity in the pedigree, we searched for homozygous by descent variants, i.e. we selected homozygous variants shared by the three affected individuals (F1-VII.10, F1-VII.16 and F1-VIII.12), which were heterozygous in both unaffected parents (F1-VI.3 and F1-VI.4). To refine the obtained lists of candidates, filtering was performed by removing all variants with a frequency above 1% on the Genome Aggregation Database (gnomAD, http://gnomad.broadinstitute.org/). Additional filtering was then performed using several frequency datasets: Iranome (https://iranome.com/) (Fattahi, Beheshtian et al. 2019), the Greater Middle East (GME) Variome (http://igm.ucsd.edu/gme/) (Scott, Halees et al. 2016), and an *in-house* exome database. After filtering using the threshold of 1%, the list of variants is refined by removing variants either with Minor Allele Frequency (MAF) above 1% in subpopulations in the databases or which have at least one homozygous individual in the database.

To predict the deleterious effect of the identified sequence variations, different bioinformatics tools were applied: MutationTaster (Schwarz, Rödelsperger et al. 2010), SIFT (Kumar, Henikoff et al. 2009), PolyPhen-2 (Adzhubei, Schmidt et al. 2010), UMD predictor (Salgado, Desvignes et al. 2016) and CADD (Kircher, Witten et al. 2014). The effect of variants on splicing was evaluated using SpliceAI (https://spliceailookup.broadinstitute.org/)(Jaganathan, Kyriazopoulou Panagiotopoulou et al. 2019) and Human Splicing Finder (Desmet, Hamroun et al. 2009)(https://genomnis.com/hsf).

##### Families 2 and 3

For Families 2 and 3, whole genome sequencing was performed on a trio in each family (Patient F2-II.2 and her parents F2-I.1 and F2-I.2 in Family 2, Patient F3-IV.2 and her parents F3-III.1 and F3-III.2 in Family 3) through the 2025 French Genomic Medicine Initiative (Plan France Médecine Génomique 2025–PFMG2025), on the SeqOIA and the AURAGEN platforms respectively.

Briefly, libraries were prepared using the Illumina TruSeq DNA Polymerase Chain Reaction-Free Sample Preparation kit (Illumina Inc., San Diego, CA, USA) and then sequenced on a NovaSeq6000,or NovaSeq X+ instrument (Illumina Inc., San Diego, CA, USA), with ae 150 bp paired-end protocol. The analysis of WGS was performed as previously described for germline analyses(contributors 2025). Raw data were mapped to the built of the human genome (GRCh38) using BWA-MEM v0.7.17 (Li and Durbin 2009, Li 2013). Variant calling was subsequently performed using GATK v4.1.8.0 (McKenna, Hanna et al. 2010) and annotation was done using VEP v98.3. Variants were then presented to the interpreting practitioners through in-house web-interfaces (gLeaves :https://gleaves.laboratoire-seqoia.fr/ or cutevariant : https://cutevariant .labsquare.org/). Variants were classified according to the ACMG-AMP guidelines, notably with the help of the mobidetails annotation platform(Baux, Van Goethem et al. 2021) (https://mobidetails.chu-montpellier.fr/).

##### Family 4 and 5

For Family 4 and 5, we re-analyzed WES data from a cohort of 76 Italian patients diagnosed with CMT and an ethnically-diverse cohort of over 20,000 patients affected by neurological disease part of the Koios database, and whose initial analyses had yielded no pathogenic findings. WES had been performed on the probands (F4-II.1; F5-II-1) of the two families. Exome capture was conducted using the Agilent SureSelect Human All Exon V6 kit, and sequencing was carried out on an Illumina HiSeq 4000 platform. Reads were aligned to the GRCh38 human reference genome using BWA-MEM v0.7.17. Variant calling was performed using DeepVariant, and variants were annotated and filtered using VEP and the platform VarAFT. Variants were prioritized based on homozygosity consistent with autosomal recessive inheritance, absence or extreme rarity in public population databases (gnomAD), and deleterious effect predictions supported by multiple in silico tools.

#### Segregation Analysis by capillary Sanger Sequencing

Candidate variants identified by WES in patients were validated by Sanger sequencing of PCR-amplified fragments in available family members. DNA available for family 1: controls: VII.1, VII.3, VII.8, VII.14, VIII.3, VIII.17, VIII.20; healthy carriers (heterozygous): VI.3, VI.4, VII.2, VII.5, VII.12, VIII.9, VIII.10, VIII.11, VIII.18, VIII.19, IX.1, IX.2; patients (homozygous): VII.7, VII.10, VII.11, VII.16, VIII.12, VIII.13. DNA available for family 2: patient (compound heterozygous II.1). DNA available for family 3: patient (homozygous II.1). The primers used are the following: Kctd11-human-F: 5’-CCGGCACATCCTCAATTTCC-3’; Kctd11-human-R: 5’-AGAGACCTGGAGTTGAGCAG-3’. PCR products were purified by mixing with an equal volume of AMPure beads (Beckman Coulter, USA) according to the manufacturer’s instructions. Both strands were sequenced as described in Jobling *et al*. (Jobling, Assoum et al. 2015). Electrophoregrams were aligned to the reference sequence using Sequencher v5.4.6 (Genecodes, USA).

### Over expression analysis

#### Plasmids

The pCMV-Myc-lKCTD11 (KCTD11 long isoform: NM_001363642) WT and mutant pCMV-Myc-lKCTD11c.493del (NM_001363642) constructs were obtained from Vector Builder (USA). The Myc tag sequence used was: 5’-ATGGAACAAAAACTCATCTCAGAAGAGGATCTG-3’ was cloned at the 5’ end of the KCTD11 coding sequence. Plasmids pCMV-Myc-lKCTD11c.601del and pCMV-Myc-lKCTD11c.217C>T were created through site-directed mutagenesis using the pCMV-Myc-lKCTD11 WT construct as a template. Mutagenesis reactions were performed using the Quikchange II XL Site Directed Mutagenesis kit (#200521, Agilent Technologies, USA), following the manufacturer’s instructions, and using the following primers: pCMV-Myc-lKCTD11c.601delC forward 5’-GAGTGGGCCCCCGCCCCGTGGAAC-3’ and reverse 5’-GTTCCACGGGGCGGGGGCCCACTC-3’; pCMV-Myc-lKCTD11c.217C>T: forward: 5’-AAATTGAGGATGTGCCAGAAGGCCTTGCCATCC-3’ and reverse: 5’-GGATGGCAAGGCCTTCTGGCACATCCTCAATTT-3’; pCMV-Myc-lKCTD11c.310C>T: forward: 5’-GAGGGGCCGGATCTAGTAGAAGTCAGCCT-3’ and reverse: 5’-AGGCTGACTTCTACTAGATCCGGCCCCTC-3’. Plasmid DNA was purified using the NucleoBond Xtra Midi Plus EF kit (Macherey-Nagel, Germany), designed for endotoxin-free plasmid DNA purification. Successful mutagenesis, integrity of the open reading frame (ORF), and CMV promoter sequence were verified by DNA sequencing.

#### Transfection

Plasmid DNA encoding WT or mutated lKCTD11 was introduced into HEK293 cells using Lipofectamine 3000 (Thermo Fisher Scientific, USA), following the manufacturer’s protocol. Briefly, 1,500 ng of plasmid DNA was mixed with 1.8 µL of Lipofectamine in Opti-MEM medium and incubated for 15 min at room temperature before being added to the cells. Cells were incubated for 48 h post-transfection before cell harvesting. For autophagy inhibition experiments, cells were incubated with 200 nM Bafilomycin A1 or vehicle control (DMSO) for 6 hours. After treatment, cells were harvested for protein extraction and subsequent Western blot analyses.

### Cell culture

#### Primary cell culture

##### Dorsal root ganglia neurons/Schwann cells (DRGN/SC) co-cultures

Mouse dorsal root ganglia (DRG) were isolated after dissecting the spinal cords of embryonic day 13.5 (E13.5) embryos and seeded onto 12 mm glass coverslips, as previously described (Poitelon and Feltri, 2018). In detail, after isolation, DRGs were dissociated and incubated in 0.25% Trypsin solution (#25200056, Thermo Fisher Scientific, USA) for 45 minutes at 37°C. DRGs were then mechanically dissociated, and approximately 50,000 cells were counted using an automated cell counter (Countess 3, Thermo Fisher Scientific, USA) and seeded onto Matrigel (#356234, Corning, USA)-coated coverslips in C-medium, composed of MEM medium (#11090081, Thermo Fisher Scientific, USA), 2 mM L-glutamine (#25030024, Thermo Fisher Scientific, USA), 10% FBS (#10270098, Thermo Fisher Scientific, USA), 4 mg/ml D-glucose (#G5146, Sigma-Aldrich, USA), and 50 ng/ml NGF (#N6009, Sigma-Aldrich, USA). After 24 hours, the C-medium was replaced with NB-medium, composed of Neurobasal medium (#21103049, Thermo Fisher Scientific, USA), 4 g/l D-glucose (#G5146, Sigma-Aldrich, USA), 2 mM L-glutamine (#25030024, Thermo Fisher Scientific, USA), 50 ng/ml NGF (#N6009, Sigma-Aldrich, USA), and 1X B27 supplement (50X stock, #17504044, Thermo Fisher Scientific, USA). The NB-medium was changed every two days. After 5 days in NB-medium, myelination was induced by adding 50 µg/ml ascorbic acid (#A0278, Sigma-Aldrich, USA) to the C-medium, for either 8 days (reaching D14 of total culture) or 16 days (reaching D22 of total culture).

##### Dorsal root ganglia neurons culture

Dorsal root ganglia (DRG) neurons were isolated and dissociated as described in the previous section. Approximately 50,000 cells were plated per well. One day after plating, cultures were maintained for 8 days in alternating culture media to eliminate non-neuronal cells. Specifically, cultures were alternately incubated with NB-medium (previously described) and NB-medium supplemented with a combination of 10 µM 5-fluoro-2’-deoxyuridine (FdU; #F-0503, Sigma, USA) combined with 10 µM uridine (#U-3003, Sigma, USA). Each medium condition was applied for two-day intervals over the 8-day period.

##### Schwann cell culture

Primary rat Schwann cells (SCs) were prepared from the sciatic nerves of postnatal day 3 pups. SCs were maintained in Dulbecco’s Modified Eagle’s Medium (DMEM; #41965039, Thermo Fisher Scientific), supplemented with 2 mM L-glutamine (#25030024, Thermo Fisher Scientific), 10% fetal bovine serum (FBS; #10270098, Thermo Fisher Scientific), 2 μM forskolin (#344270, Merck), and 2 ng/ml recombinant human NRG1-β1 (#396-HB, R&D Systems), as described in (Poitelon and Feltri 2018), SCs were not used beyond the fourth passage.

#### HEK293 cell line

HEK293 (Human embryonic kidney, #CRL-1573) line was provided by ATCC (Manassas, Virginia, USA). HEK293 cells were cultured in DMEM (#41965, Thermo Fisher Scientific, USA) supplemented with 10% of FBS (#15000-97036, Thermo Fisher Scientific, USA) and 1% of penicillin/streptomycin (#15070063, Thermo Fisher Scientific, USA).

### Animal studies

#### Mouse model

In this study, we used the knockout mutant of the potassium channel tetramerization domain containing 11 gene (*Kctd11^-/-^*; C57BL/6NJ-Kctd11em1(IMPC)J/Mmjax (MMRRC Strain #046124-JAX]), generated by CRISPR-Cas9 in the frame of the Knockout Mouse Phenotyping Program (KOMP2) at The Jackson Laboratory. The genetic modification led to the deletion of 697 base pairs in exon 1 of the *Kctd11* gene, causing a change in the amino acid sequence after residue four and resulting in a truncation 103 amino acids later, followed by read-through into the 3’ UTR. Phenotypic characterization of this strain, including morphological and compartmental analyses, was not reported at the start of the project. Now, the phenotypic characterization is reported by the International Mouse Phenotyping Consortium (https://www.mousephenotype.org/data/genes/MGI:2448712#data) The effectiveness of the knock-out was verified by bulk mRNA sequencing, which confirmed that *Kctd11* is no longer expressed, as no reads were detected in the *Kctd11^-/-^* condition. The presence of the deletion was confirmed by PCR genotyping using the following primers: Kctd11-ms-F (5’-TCCCGTGACCCTAAATGTGG-3’) and Kctd11-ms-R (5’-GAGGTTTTAGAAGAGCCACAGG-3’). The expected amplicon size is 150 bp for the *Kctd11^-/-^* allele and 846 bp for the WT allele. Animals were housed in an animal facility with a 12 h light/12 h dark environment and *ad libitum* access to water and a standard diet. All experiments were done in accordance with a national appointed ethical committee for animal experimentation (Ministère de l’Education Nationale, de l’Enseignement Supérieur et de la Recherche; Authorization No. #46895-2024011716006728 v2).

#### Behavioral tests

To assess locomotor performances of our knock-out *Kctd11^-/-^* mice, over age, we performed a longitudinal study, using the three following tests : rotarod, grip strength and gait analysis (Brooks and Dunnett 2009). Each test was repeated on the same set of animals at different ages (6-12 and 18 months). We used three groups (*Kctd11^-/-^* (KO), *Kctd11^+/-^*(Het) and WT) of sex and age matched, littermate animals at 6 (n=18 *Kctd11^-/-^*, n=16 Het and n=16 WT) and, 12 (n=18 *Kctd11^-/-^*, n=16 Het and n=16 WT) and 18 (n=16 *Kctd11^-/-^*, n=14 Het and n=14 WT) months old.

Locomotion was evaluated by the rotarod test. Motor coordination and balance were evaluated using an accelerating rotarod apparatus (TSE systems). The protocol consisted of three phases over 3 days as described in (Bernard-Marissal, van Hameren et al. 2019): i) Day 1 (Habituation): Mice were placed individually on the stationary rotarod to freely explore the apparatus for 5 minutes to minimize stress and familiarize them with the device. ii)Day 2 (Training): Each mouse underwent three timed trials on the rotarod, with the rod accelerating from 4 rpm to 40 rpm over 5 minutes. The latency to fall (time in seconds before the mouse fell off or completed a full rotation while hanging on) was recorded automatically. A rest period of at least 15 minutes was allowed between trials. ii) Day 3 (Evaluation): After one day of rest, mice were tested again with three timed trials under the same acceleration conditions as during training. The maximal latency to fall across the three trials was calculated for each mouse and used for subsequent analysis.

The grip test was used to quantify grip strength, considered a reliable indicator of neuromuscular function in mice. The test was carried out in a single day, in two stages. For habituation, mice were first placed in the experimental room and left to rest for at least 30 minutes, to minimize the effects of stress or environmental change. For the measurements, each mouse was gently grasped by the base of the tail, then brought up to the grip plate so that the front paws (or all paws, depending on the condition being tested) could grip the metal grid. Once the animal was correctly positioned, its body was held in a quasi-horizontal posture, then slowly pulled backwards in the horizontal plane, in line with the load cell, until it released its grip. In this way, the maximum force exerted by the mouse at the moment of traction was automatically recorded by the measurement system. For both conditions, front paws only or all paws, five trials were performed per mouse, with a five-minute rest interval between each measurement to limit fatigue or learning effects. Finally, the analyses were performed on the three best values, in accordance with standard practice.

For the gait experiment, gait was analyzed during spontaneous walk using an automated gait analysis system (Gaitlab, Viewpoint, France). Before recording footprints, mice were acclimated and trained to walk on the Gait system for two days. During the testing session, a minimum of 2-3 completed runs were collected. We focused the analysis on intensity-based parameters, paw-size as well as gait/posture, as previously described in(El-Bazzal, Rihan et al. 2019). Animals that did not complete at least 2 successful runs (stalling or reversing during gait,…) were removed from the study.

#### Nerve Conduction Velocities measurements

Nerve conduction velocities (NCVs) were measured in *Kctd11^-/-^*, *Kctd11^+/-^* and WT mice at 12 (n=5 *Kctd11^-/-^*, n=5 Het and n=5 WT) or 18 months (n=5 *Kctd11^-/-^*, n=4 Het and n=5 WT), on two separate groups of animals, as described (Schulz, Walther et al. 2014). Briefly, mice were anesthetized by isoflurane inhalation (5% induction in the induction cage, followed by 1.5% maintenance by mask), with a gas mixture consisting of oxygen-enriched air. Throughout recording, mice were placed on a heated platform to maintain a stable body temperature. NCVs were measured using the PowerLab system coupled to LabChart software (ADInstruments). We used five electrodes inserted under the skin without incision: two recording electrodes were positioned at the base of the hind paw: one subcutaneous, the other intramuscular, two stimulation electrodes were placed either proximally (near the sciatic node and at the base of the tail) or distally (at the ankle and at the base of the gastrocnemius muscle). Stimulation and detection times were in the millisecond range. Three measurements were taken per paw, for a total of six per animal, in order to limit artifacts and guarantee data reproducibility. The total duration of the procedure was approximately 10 to 15 minutes for each animal. At the end of the recording, the animals were returned to their experimental cage, positioned under a red-light heat lamp, and kept warm until they are fully awake. They were then placed temporarily in a recovery cage, under supervision, for around ten minutes. Once fully awake, the animals were returned to their original cage with their conspecifics, and monitored post-procedure using a standardized assessment grid.

The recorded data were analyzed using LabChart software, enabling extraction of the following parameters: mean peak latency, mean duration of the evoked signal.

### Electron microscopy and morphometric analyses

The sciatic, tibial and saphenous nerves were dissected and fixed in 2% Paraformaldehyde (PFA) and 2.5% Glutaraldehyde in 0.1M cacodylate buffer for 5 hours. The next day, the nerves were washed three times in 0.1M cacodylate buffer and post-fixed in buffered 1% OsO4 for one hour. After washes in distilled water, the samples were contrasted in aqueous 1% uranyl acetate. Samples were then dehydrated in graded series of ethanol baths (30 minutes each) and infiltrated with epon resin in ethanol (1:3, 2:2, 3:1) for 2 hours for each, and finally in pure resin overnight. The next day the nerves were embedded in fresh pure epon resin and cured for 48h at 60°C. 500 nm semi-thin and 70 nm ultra-thin sections were performed on a Leica UCT Ultramicrotome (Leica, Austria). Semi-thin sections were stained with toluidine blue and ultrathin sections were deposited on formvar-coated slot grids. The grids were contrasted using uranyl acetate (10 minutes) and lead citrate (5 minutes) and observed using an FEI Tecnai G2 at 200 KeV. The acquisition was performed on a Veleta camera (Olympus, Japan). The proportion of fibers having out- and infoldings was counted on ten fields corresponding to a range of 600-800 fibers for each sciatic nerve of at least three animals. To perform g-ratio analysis, digitalized images of fiber semithin sections of the sciatic nerves were obtained with a 100× objective of a phase-contrast microscope (BX59, Olympus). At least ten images from three different animals per genotype at 3-, 6-, 12- and 18-month-old were acquired. he g-ratio was calculated as the ratio of the mean axon diameter (without myelin) to the mean fiber diameter (axon plus myelin sheath), using MyelTracer software (Kaiser, Allen et al. 2021) available at https://github.com/HarrisonAllen/MyelTracer.

### Transcriptomic analysis

#### RNA extraction

For DRGN/SC co-cultures, total RNA was extracted from each condition, using Purelink silica membrane, anion exchange resin, spin-column kits, following the manufacturer’s instructions (Purelink RNA Minikit, #12183018A, Thermo Fisher Scientific, USA). The same procedure was performed for the rat Schwann cell samples, in triplicate. For sciatic nerve samples, immediately after dissection, nerves at P0, P5, P15, and P30 were snap-frozen in liquid nitrogen and stored at -80°C. On the day of RNA extraction, the nerves were homogenized using the FastPrep Tissue Homogenizers (MP Biomedicals, USA) in Qiazol (Qiagen). Total RNA was then extracted using RNeasy® Lipid Tissue Kit (Qiagen, #74804, Germany) as per manufacturer’s instructions.

#### Bulk mRNA-sequencing

RNA sequencing and bioinformatics analysis were performed by the Genomics and Bioinformatics facility (GBiM) of the U1251/Marseille Medical Genetics Laboratory. We performed bulk mRNA sequencing (mRNA-Seq) experiments, in triplicate or quadruplicate, using total RNA samples extracted from: i) knockout (*Kctd11^−/−^)* or WT DRGN/SC co-cultures (quadruplicates for each genotype and each time point; 16 samples total), ii) rat Schwann cells at P4 (triplicate), and iii) sciatic nerves from knockout (*Kctd11^−/−^)* or WT mice at different postnatal ages (P0, P5, P15, and P30) (quadruplicates; 32 samples total). Before sequencing, the quality of total RNA samples was assessed using the Agilent Bioanalyzer (Agilent Technologies, USA): only RNAs with RNA Integrity Numbers (RIN) above 8.5 were deemed suitable for sequencing and used for library preparation. RNA concentration was then measured using the Qubit™. For each sample, a stranded mRNA library was prepared from 300 ng of total RNA after capture of polyadenylated RNA species using the KAPA mRNA HyperPrep Kit (#KR1352-v7.21, Roche, Switzerland), following the manufacturer’s instructions. The quality and profiles of individual libraries were quantified and visualized using Qubit™ and the Agilent Bioanalyzer dsDNA High Sensitivity Kit (Agilent Technologies, USA), respectively. Indexed libraries were pooled and sequenced (2 × 100 bp paired-end) on an Illumina NovaSeq 500 platform (Illumina, USA).

#### Data processing and differential gene expression analysis

The quality of sequencing reads was assessed using FastQC v0.11.5 (https://www.bioinformatics.babraham.ac.uk/projects/fastqc/). Raw sequencing reads were mapped to the mouse reference genome (Mus musculus) genome assembly GRCm39/mm39 using STAR v2.7.2b (Dobin and Gingeras 2015) and BAM files were sorted and indexed using samtools v1.7 (Danecek, Bonfield et al. 2021). After mapping, the number of reads per feature (GENCODE vM34 annotations) was determined using Stringtie v2.1.6 (Pertea, Pertea et al. 2015). Differential gene expression analysis was performed using a Wald test with the DESeq2 v1.34.0 package (Love, Huber et al. 2014). *P*-values were adjusted for multiple testing using the method described by Benjamini and Hochberg (BH) (Benjamini et al. 1995 https://www.jstor.org/stable/2346101). Relative expressions of the most variable features between samples have been plotted as heatmaps using the Pheatmap R package, representing the Log2FC (log2 of the expression fold change) and the adjusted p-value, prepared using the EnhancedVolcano R package. Enrichment (Gene Set Enrichment Analysis, GSEA) and over-representation analysis (Singular Enrichment Analysis, SEA) of Gene Ontology (GO) terms among DEGs were performed using Mann–Whitney and hypergeometric tests, respectively, via the enrichGO function of the clusterProfiler R package (v3.10.15). For functional GO annotation, DEGs with a statistically significant adjusted p-value (Padj) < 0.01 (SEA) were selected, and each gene was assigned to GO terms related to Cellular Component (CC), Biological Process (BP), or Molecular Function (MF).

### Molecular analysis

#### Protein extraction and immunoblotting

DRGN/SC co-cultures, HEK293 cells, rat primary Schwann cells, and nerve samples were lysed in RIPA buffer supplemented with a protease and phosphatase inhibitor cocktail (#78442, Thermo Fisher Scientific, USA). Lysates were homogenized using the FastPrep-24 5G system (MP Biomedicals, USA) at 6 m/s for 40 seconds, followed by sonication with the Bioruptor VCD-200 (Diagenode, Belgium). After centrifugation at 16,000 × g for 20 min at 4°C, the supernatant was collected, and protein concentration was determined using a bicinchoninic acid (BCA) assay (#B9643-1L, Sigma-Aldrich, USA) in combination with a copper (II) sulfate solution (#C2284-25ml, Sigma-Aldrich, USA), according to the manufacturer’s instructions.

A total of 40 μg of protein per sample was loaded onto a precast NuPage 4-12% Bis-Tris gel (Thermo Fisher Scientific, USA) and transferred onto a nitrocellulose membrane (GE Healthcare, USA). TMembranes were blocked with Intercept Blocking Buffer (Li-Cor, USA) and incubated overnight at 4°C with the following primary antibodies: rabbit anti-HDAC1 (1:500, GeneTex, #GTX100513), rabbit anti-β-catenin (1:1000, Cell Signaling, #9587), rabbit anti-phospho-β-catenin (1:1000, Cell Signaling, #CST9565), mouse anti-GAPDH (1:2000, Abcam, #ab125247), mouse anti-α-tubulin (1:4000, Sigma-Aldrich, #T6074), rabbit anti-mTOR (7C10) (1:1000) (#2983T, Cell Signaling Technology, USA), rabbit anti-phospho-S6 ribosomal (Ser235/236, 1:1000, Cell signaling, #2211), rabbit anti-S6 ribosomal (1:1000, Cell signaling, #2217), rabbit anti-phospho-4E-BP1 (1:1000, Cell signaling, #2855), and rabbit anti-4E-BP1 (1:1000, Cell signaling, #9452). Following three washes with PBS containing 0.1% Tween-20 (PBS-T), membranes were incubated with secondary antibodies: IRDye® 800CW Donkey anti-Mouse IgG H&L and IRDye® 680RD Donkey anti-Rabbit IgG H&L (Li-Cor Biosciences, USA), both diluted at 1:10,000. Detection was performed using the ChemiDoc imaging system (Bio-Rad, USA). Band intensities were analyzed using the Gel Analyzer tool in ImageJ (ImageJ Software, National Institutes of Health, Bethesda, MD, USA, http://rsb.info.nih.gov/ij). For each sample, the relative protein intensity was calculated by normalizing the target protein intensity to the corresponding loading control. Data were then normalized to the control condition.

#### RT-PCR

For reverse transcription (RT)-PCR analysis, total RNA was isolated from transfected HEK293 cells or freshly dissected muscle, brain, cerebellum, sciatic nerve and spinal cord using Purelink silica membrane, anion exchange resin, spin-column kits, following the manufacturer’s instructions (PureLink RNA Mini Kit, #12183018A, Thermo Fisher Scientific, USA). cDNAs were synthesized with MultiScript^TM^ reverse transcriptase (#4368814 Thermo Fisher, USA) using random primers. Specific PCR primer sets were designed according to mRNA sequences. The following primer pairs were used for mRNA-based RT-PCR analysis: mouse Kctd11 transcripts F: 5’-GGTACGGAGAAACTGCCCTT-3’ and R: 5’-TGCAGCCCCAGTCTTTGATA-3’; human Myc-tagged KCTD11 transcripts: F: 5’-GATCTGTCTCCTCCTCCTGTG-3’ and R: 5’-AGAGACCTGGAGTTGAGCAG-3’; human ACTB transcripts: Ex4-F: 5’-CCCTGGAGAAGAGCTACGAG-3’ and Ex5-R: 5’-TCTCCTTCTGCATCCTGTCG-3’.

### Fluorescent immunolabeling

#### Fluorescent immunocytochemistry

Conditions for immunostaining of DRGN/SC co-cultures were described previously (El-Bazzal, Ghata et al. 2023), and those for hiPSC-MN and HEK293 cells were detailed in(El-Bazzal, Rihan et al. 2019). Primary antibodies used for the DRGN/SC co-cultures myelin segment quantification included: chicken anti-neurofilament NF-M (1:1000, #822701, BioLegend, USA) and rat anti-MBP (1:300, #MAB386, Merck-Millipore, Germany). For hiPSC-MN cultures, the following primary antibodies were used: mouse anti-HB9 (1:50, #81.5c10, DSHB), chicken anti-neurofilament NF-M (1:1000, #822701, BioLegend, USA), and rabbit anti-KCTD11 (1:500, #AP21901a, Abgent). For HEK293 cells, primary antibody used was: anti-c-Myc (1:500, #2276, BioLegend, USA). After two washes in 1× PBS, cells were incubated with the following secondary antibodies: for the DRGN/SC co-cultures, donkey anti-rat IgG H&L (1:1000, #150154, Alexa Fluor 555, Abcam, UK) and goat anti-chicken IgY H&L (1:1000, #A21449, Alexa Fluor 647, Invitrogen, USA); for the hiPSC-MN, donkey anti-rat IgG H&L (1:1000, #150154, Alexa Fluor 555, Abcam, UK), goat anti-chicken IgY H&L (1:1000, #A21449, Alexa Fluor 647, Invitrogen, USA), and donkey anti-rabbit IgG H&L (1:1000, #150073, Alexa Fluor 488, Abcam, UK). Coverslips were then rinsed twice in 1× PBS and mounted with Duolink mounting medium (#DU082040, Sigma-Aldrich, USA) for microscope analysis. Images were acquired using a Zeiss ApoTome.2 microscope equipped with an AxioCam MRm camera (Zeiss, Germany), an Axio Observer.Z1/7 with a Hamamatsu camera (Zeiss, Germany), and an Axio Scan.Z1 microscope equipped with a Hitachi HV-F202SCL camera (Zeiss, Germany). Image acquisition and merging were performed with ZEN software (Zeiss, Germany), and subsequent processing was done using ImageJ software.

#### Immunohistochemistry

Sciatic nerves were collected and fixed for eight hours in 4% paraformaldehyde (PFA) at room temperature (RT), washed with 1× PBS, infiltrated overnight (O/N) in 20% sucrose at 4°C, embedded in OCT compound (#TFM-5, Microm Microtech, France), and snap-frozen in liquid nitrogen. Sciatic nerve cross-sections (30-μm thick) were prepared and left to air dry for 2 hours at RT. Sections were then rehydrated in 1× PBS for 10 minutes, followed by permeabilization with 0.2% Triton X-100 in PBS for 15 minutes at RT. Blocking was performed for 1 hour at RT in a solution containing 10% fetal bovine serum, 2% bovine serum albumin, and 0.2% Triton X-100 in 1× PBS. Primary antibody incubation was carried out overnight at 4°C using the following antibodies: rabbit anti-KCTD11 (1:500, #AP21901a, Abgent), chicken anti-NF-M (1:1000, #822701, BioLegend, USA), and rat anti-MBP (1:300, #MAB386, Merck-Millipore, Germany). After two washes in 1× PBS, sections were incubated with secondary antibodies for 2 hours at 4°C: donkey anti-rat IgG H&L (1:1000, #150154, Alexa Fluor 555, Abcam, UK), goat anti-chicken IgY H&L (1:1000, #A21449, Alexa Fluor 647, Invitrogen, USA), and donkey anti-rabbit IgG H&L (1:1000, #150073, Alexa Fluor 488, Abcam, UK). Finally, slides were mounted using Prolong™ Diamond Antifade Mountant with DAPI (#P36962, Life Technologies, USA). Z-stack images were acquired using an Axio Observer.Z1/7 microscope equipped with a Hamamatsu camera (Zeiss, Germany).

#### Analysis of myelination

To quantify myelination under each condition, 15 fields per coverslip were acquired following the same predefined trajectory using a Zeiss ApoTome.2 microscope (Zeiss) with a ×20 objective. The number of myelin segments, visualized as myelin basic protein (MBP)-positive segments, was counted in each coverslip for both WT and *Kctd11^-/-^* conditions. For length measurements (provided as supplementary data), myelin segment lengths were measured using the NeuroJ plugin in ImageJ.

### Statistical analyses

Statistical analyses were performed using an unpaired two-tailed Student’s t-test, one-way ANOVA, or two-way ANOVA for parametric data, as appropriate. The specific statistical test used is indicated in each figure legend. Data are presented as mean ± SEM. Statistical significance was set at P < 0.05 (*P < 0.05, **P < 0.01, *P < 0.001).

### 3D protein folding analysis

The three-dimensional (3D) structures of WT and mutated long-KCTD11 proteins were predicted using the AlphaFold2 Colab notebook (Jumper, Evans et al. 2021). Protein sequences were provided as input, and structural predictions were generated using the default parameters. The highest-ranked models, based on the predicted local distance difference test (pLDDT) scores, were selected. The predicted structures were then visualized and analyzed using ChimeraX (UCSF ChimeraX, USA; (Goddard, Huang et al. 2018)).

### Data availability

The authors confirm that the data supporting the findings of this study are available within the article and/or its Supplementary material. These data are available from the corresponding author, upon reasonable request.

## Results

### Clinical findings

10 patients from 5 unrelated families of diverse ethnic background (Palestinian, French, Italian and Egyptian) are included in this study. Clinical characteristics of all patients included in this study are summarized in Table 1.

**Table 1.**
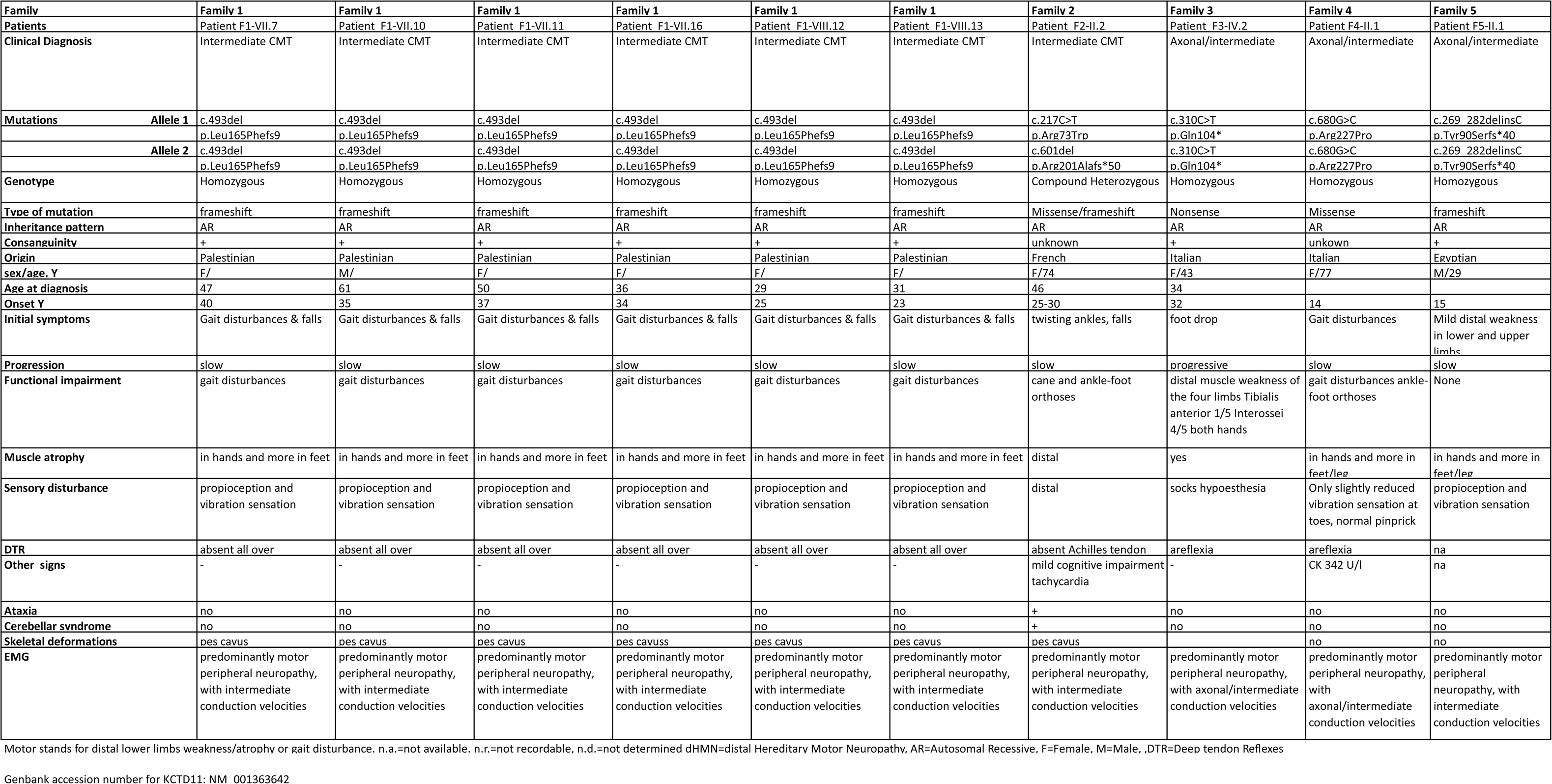
Genetic, clinical and electrophysiology findings of all patients described in the study.

### Family 1

Family 1 is a large Palestinian inbred family from Lebanon with six individuals affected by autosomal recessive intermediate form of Charcot-Marie-Tooth diseases (RI-CMT). Patients F1-VII.7, F1-VII.10, F1-VII.11, F1-VII.16, F1-VIII.12 and F1-VIII.13 had a clinical examination. The affected individuals (F1-VII.7, F1-VII.10, F1-VII.11, F1-VII.16, F1-VIII.12, F1-VIII.13) presented with a moderate sensorimotor impairment predominantly affecting the lower limbs distally. The disease occurred in the mid-thirties/ early-forties and its course was slowly progressive. Muscle weakness and atrophy were mildly to moderately present in the hands and mainly the feet., except the patient F1-VII.7, who had severe amyotrophy and weakness in both hands and legs with marked steppage gait. There was an involvement of the proprioception and vibration sensation modalities in the feet, in all patients Feet and hand deformities were variable: most patients had mild pes cavus deformities and very discrete hand deformities. The oldest patient (F1-VII-10, age 84 years) required a cane for ambulation. The deep tendon reflexes were all abolished in 4 limbs, in all affected patients Electrophysiological studies were performed for patients F1-VII.10, F1VII.11, F1-VII.16, F1-VIII.12 and F1-VIII.13 (ages 35-65 years) and unaffected individuals F1-VI.1, V, F1-VI.5, F1VII.2 and F1-VII.8 (ages 60-80). Results shown in Table 2 for electrophysiological measurements performed in patient F1-VIII.13 at the age of 39 years revealed severe drop in the CMAPs and slowing in the motor and sensory nerve conduction velocities in the nerves of her lower limbs. MNCVs values at the median nerve were reduced to values below45 m/s and compatible with intermediate CMT disease as defined by Berciano et al (Berciano, Garcia et al. 2017) (MNCVs values between 25 and 45 m/s at the median nerve with normal amplitudes).

**Table 2.**
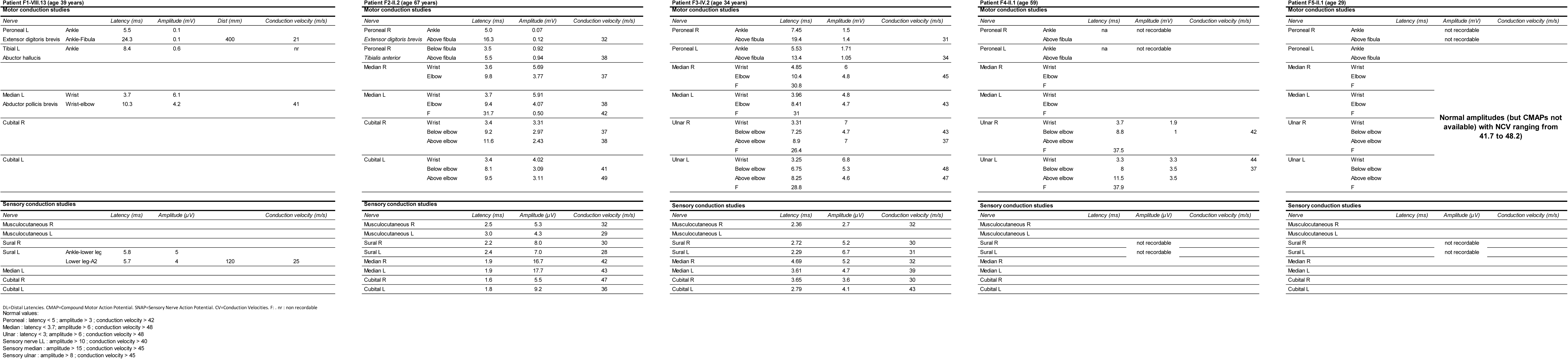
Nerve conduction studies in patients from family 1, family, family 3, family 4 and family 5.

### Family 2

Patient F2-II.2, a 74-year-old woman, was born to unrelated parents. She has a sister with gait disturbances who required a wheelchair at the age of 78, a brother (whose EMG was normal) who died at age 50 from lung cancer, and two children aged 43 and 47 who are in good health and have normal EMG results. She did not exhibit delayed motor development or motor problems during childhood. Her first symptoms appeared between the ages of 25 and 30, including ankle twisting, falls, and high-arched feet (pes cavus). A diagnosis of Charcot-Marie-Tooth disease (CMT) was made at age 46, after which she was gradually lost to follow-up. Clinical deterioration was noted around age 52, with worsening of gait disturbances and the onset of a cerebellar syndrome and dysarthria. On clinical examination at age 67, she presented with distal sensorimotor amyotrophic weakness, predominantly in the lower limbs, with bilateral foot drop. Muscle strength scores (MRC scale) were 4/5 for finger extensors and flexors, 3/5 for interossei, and 2/5 for anterior tibialis, gastrocnemius, and peroneal muscles. Achilles tendon reflexes were absent. Central nervous system findings included dysmetria, dysarthria, and intention tremor.

At her most recent clinical examination at age 74, she was walking with a cane and ankle-foot orthoses. A neuropsychological assessment showed attention difficulties characterized by fluctuations, as well as impaired working memory processes (both maintenance and manipulation). Executive function testing showed difficulties in mental flexibility and a slight ideational slowing, with persistent problems in lexical-semantic access and selection. Episodic memory (verbal and visual) performance was slightly reduced but within the normal range, with retrieval being the main issue.

Cardiological examination revealed longstanding tachycardia, but echocardiography was normal. The last EMG performed at the age of 67 showed a predominantly motor peripheral neuropathy, with intermediate conduction velocities and poorly excitable nerves, but with little progression. Brain MRI was normal. Spinal MRI showed cervical and lumbar disc disease. Lumbar puncture, as well as visual and auditory evoked potentials, were normal. From a metabolic perspective, tests including transferrin isofocusing, pristanic and phytanic acids, vitamin E, homocysteine, arylsulfatase, very long-chain fatty acids, acanthocytes, and lipid panel were all normal.

Muscle biopsy showed a few COX-deficient fibers. The mitochondrial respiratory chain study was normal.

From a genetic standpoint, testing for *MERRF*, *MELAS*, N*ARP* mutations was negative. NGS panels for CMT, cerebellar ataxias, and mitochondrial diseases were also negative, as was the sequencing of the *CYP27A1* gene (associated with cerebrotendinous xanthomatosis) and the testing for CANVAS.

### Family 3

Patients F3-IV.2 is a 43 years old French female born to consanguineous parents, originating from Calabria (Italy). After a fracture of the right 5th metatarsal, at the age of 32, the patient noticed, that she had difficulty walking with bilateral steppage. Clinical examination revealed a motor deficit of the tibialis anterior predominantly on the right side (MRC 1/5 vs 2/5) with muscle atrophy, sock hypoesthesia and diffuse tendon areflexia. The patient did not report any delay in motor acquisition or scoliosis in childhood. Her brother, sister, daughter, and parents have no neurological deficits. The Electrodiagnostic assessment (EDX), performed at age 34 shows a length-dependent intermediate motor and sensory neuropathy within the intermediate range (25-45 m/s at the median nerve) as defined by Berciano et al (Berciano, Garcia et al. 2017).

After ten years of follow-up, the motor deficits progressively increased. The tibialis anterior is evaluated as 1/5 bilaterally and the interosseous muscles of both hands as 4/5. The patient walks with two anti-step orthoses.

### Family 4

Patient F4-II.1 is a 76-year-old Italian female, born to parents from two neighboring small towns in Italy. There was no reported delay in motor milestones during childhood. Symptoms began in adolescence with very slowly progressive distal weakness, initially affecting the lower limbs. By her early 50s, she developed increasing difficulty with fine motor tasks in the hands, such as buttoning. Ankle-foot orthoses were prescribed in her mid-60s but were never used. Clinical examination was performed at the age of 70 and revealed bilateral steppage gait and complete foot drop (MRC grade 0 for both plantarflexion and dorsiflexion). In the upper limbs, she exhibited severe hand weakness, with MRC grade 1 in abductor pollicis brevis and first dorsal interosseous, and grade 4+ in extensor digitorum. Proximal muscle strength was preserved, and cranial nerve examination was unremarkable. Sensory involvement was mild, limited to reduced vibration sense at the toes, with normal pinprick sensation. Deep tendon reflexes were absent. Charcot-Marie-Tooth Examination Score (CMTES) was 12, indicating a moderate disease severity. Nerve conduction studies performed at age 59 revealed a length-dependent axonal/intermediate neuropathy, with ulnar motor conduction velocities ranging between 37 and 44 m/s.

### Family 5

Patient F5-II.1 is a 29-year-old Egyptian male, born to consanguineous parents (third cousins). There is no known family history of neuromuscular disorders. Early motor development was reported as normal. Symptoms began at age 15 with mild distal weakness in the lower limbs. The disease progression was remarkably slow, such that at the time of examination at age 29, gait was preserved without the need for AFOs, or walking aids. Nerve Conduction Studies (NCS) revealed an intermediate neuropathy in the upper limbs, with normal CMAPs and motor NCVs ranging from 41.7 to 48.2 m/s. In the lower limbs, common peroneal CMAPs and sural SAPs were absent.

### Genetic findings

More than 15 years ago, we first realized a genome wide screen in Family 1, using 400 polymorphic Single Tandem Repeat microsatellite markers with an average intermarker distance of 10 cM, We performed Homozygosity mapping (Lander and Botstein 1987) as previously described (Delague, Bareil et al. 2000), but were not able to assign a homozygous by Descent (IBD) locus in the family. At the time, this was interpreted as the result of multiple recombination events which occurred during meiosis over generations and led to a homozygous by descent region smaller than the intermarker microsatellite distance (10 cM, ∼10 Mb), and therefore non detection of the IBD region in our data. Later, whole-genome sequencing was undertaken on five individuals from the family: three affected individuals F1-VII-10, F1-VII-16 and F1-VIII-12 and two unaffected parents F1-VI-3 and F1-VI-4, with the aim of performing Identical by Descent (IBD) analysis of the WGS data, as described in the Material and Methods section.

This approach led to the identification of 9 rare (MAF<1% in gnomAD) Identical By Descent (IBD) Single Nucleotide (SNV) variants (**Supp Table 1**), located on the same apparently IBD region on chromosome 17p13.2-p12. Search for Runs of Homozygosity (ROH) using plink allowed to identify three adjacent IBD regions on chromosome17p13.2-: rs72839804-rs9303210 (938 kb), rs56394091-rs9909211 (2.59 Mb) and rs9905443-rs58082448 (1.85 Mb). This result being in contradiction with the results obtained by genome wide screen, we came back to those data and noticed that the individual F1-VII.3 had been wrongly labeled as affected in the first genome wide-screen in place of F1-VII.11. Re-analysis of these data, considering this new information, led, indeed, to the assignment of a 11.2Mb IBD region on chromosome 17p13.3.-p12 between STR markers D17S831 (AFM058XF4) and D17S799 (AFM192YH2). All genes included in this region are covered 100% at 10X in the WGS of the patients. Of the 9 IBD variants identified in the WGS, 7 were eliminated by further investigation of their MAFs in gnomAD v4.1.0 (Supp Table 1). The two remaining homozygous variants, the c.493del/p.(Leu165Phefs*9) frameshift in *KCTD11* (NM_001363642) and the synonymous c.306G>C p.(Thr102Thr) change in *SPEM2* (NM_175734), were both extremely rare (f=1.86.10^-6^ and 6.19.10^-7^ in gnomAD v 4.1.0 respectively, see Supp Table 1).We selected the homozygous frameshift variant in *KCTD11* (NM_001363642) as the only candidate variant responsible for RI-CMT in the patients, considering the fact that the variant in *SPEM2* is synonymous and not predicted to affect splicing by spliceAI (Jaganathan, Kyriazopoulou Panagiotopoulou et al. 2019) and Human Splicing Finder (Desmet, Hamroun et al. 2009), and most importantly, because it is specifically expressed in the testis and therefore not expressed in the Peripheral Nervous System. The c.493del single-base deletion in *KCTD11* (NM_001363642) induces a frameshift p.(Leu165Phefs*9), likely resulting in a truncated or absent protein and therefore loss of function.

Segregation analysis, by Sanger Sequencing, confirmed that the identified candidate variant in *KCTD11* segregates with the disease, with all six affected individuals (F1-VII.7,.10,.11,.16, -F1-VIII.12 and.13) homozygous, while obligate carriers (F1-VI.3 andVI.4) are heterozygous and other unaffected family members are either heterozygous (F1-VII.2, VII.5, VII.12, VIII.9, VIII.10, VIII.11, VIII.18, VIII.19, IX.1 andIX.2), or homozygous for the normal allele (F1-VII.1, VII.3, VII.8, VII.14, VIII.3, VIII.17 andVIII.20) (**Figure 1)**. This homozygous c.493del p.(Leu165Phefs*9) variant *KCTD11* (NM_001363642), was in consequence considered as the candidate pathogenic variant causing this particular form of RI-CMT in family 1.

Three additional patients from unrelated French (Families 2-3), Italian (Family 4) and Egyptian (Family 5) families were then identified with bi-allelic variants in *KCTD11*, further supporting the hypothesis of this gene being a new culprit gene in RI-CMT. French Family 3 is of Italian (Calabrian) descent. In Family 2, WGS identified compound heterozygous variants in patient F2-II.2 in *KCTD11*: NM_001363642:c.[601del];[217C>T], NP_001350571: [p.Arg201Alafs*50];[p.Arg73Trp] (**Figure 1**). Visual inspection of the reads on IGV allowed to demonstrate that the two variants are not on the same allele, and therefore that they are compound heterozygous. In Family 3, a homozygous nonsense c.(310C>T) p.(Gln104*) variant was identified in *KCTD11* (NM_001363642) in the WGS from patient F3-IV.2..

The patient F4-II.1 from Family 4 harbored a homozygous missense variant in *KCTD11*; c.(680G>C) (NM_001363642), p.(Arg227Pro) (NP_001350571). The Arginine at amino acid position 227 is conserved across mammals, and predicted to be damaging by multiple in silico prediction tools. Finally, in Family 5, the patient F5-II.1 harbored a homozygous variant in *KCTD11*: c.(269_282delinsC) (NM_001363642), p.(Tyr90Serfs*40) (NP_001350571), resulting in the deletion of 14 nucleotides from position 269 to 282 of the coding DNA sequence and the insertion of a single cytosine. This frameshift mutation disrupts the reading frame beginning at codon 90, replacing the native amino acid sequence with aberrant residues and introducing a premature termination codon 40 amino acids downstream.

All five variants are extremely rare or absent in the gnomAD v4.1.0 database, with no homozygous described, as detailed in **Supplementary Table 2C**, with no homozygous reported.

KCTD11 is a single-exon gene encoding two different isoforms by alternative initiation: a short 232 amino acids (aa) isoform (NM_001002914, NP_001002914) and a long 271 aa isoform (NP_001350571), encoded by the human canonical transcript (NM_001363642) and referred to as lKCTD11. The two isoforms differ only at their N-terminal extremity, where the short isoform has a shorter, incomplete POZ/BTB domain.

The p.(Leu165Phefs*9) and p.(Arg201Alafs*50) changes are frameshift variants that introduce a premature stop codon, predicted to result in a truncated or absent protein. Finally, the nonsense p.(Gln104*) variant creates a stop codon, which may also result in a truncated protein.The Arginine at amino acid (AA) position 73 is highly conserved among vertebrates and the Arginine at amino acid position 227 is conserved across mammals.

The p.(Arg73Trp) substitution occurs within the POZ/BTB domain, and is predicted by Alphafold Missense to be pathogenic. KCTD11 gene being single exon, we do not expect any effect on splicing as confimed by SpliceAI and Human Splicing Finder. However, the p.Arg227Pro missense variant is predicted deleterious by PrimateAI-3D

Given the recessive inheritance of the disease, we postulate that these variants lead to a loss-of-function effect, either by generating a non-functional protein or by activating degradation pathways that preclude protein accumulation.

Based on these results, we propose that the bi-allelic variants identified here in *KCTD11* are pathogenic and that KCTD11 is a new causative gene in CMT, more specifically responsible for autosomal recessive intermediate CMT.

### *KCTD11* mutations induce post-translational protein degradation

To further support the pathogenicity of the identified mutations, we investigated their impact on KCTD11 expression in human cells.

We first used the AlphaFold2 structural model to predict how the variants impact the three-dimensional structure of the mutants (Fig. 2A). As expected, the two frameshift variants p.(Leu165Phefs*9) (truncating 99 amino acids) and p.(Arg201Alafs*50) (truncating 22 amino acids), as well as the nonsense variant p.(Gln104*) (truncating 168 amino acids) considerably disrupted the predicted 3D structure, leading to pronounced protein misfolding. In contrast, the p.(Arg73Trp) substitution does not appear to affect the overall 3D conformation. However, this mutation is located within the highly conserved POZ/BTB domain protein, a region critical for protein stability and oligomerization of KCTD11(Correale, Pirone et al. 2011). Therefore, these structural insights suggest that the identified mutations probably compromise KCTD11 stability and function, either due to post-transcriptional or post-translational mechanisms. To investigate this, we sought to examine KCTD11 expression levels, by Western Blot, in patients’ cells from Family 1 and 2, for whom we had immortalized lymphoblastoid cells. Unfortunately, the protein was not expressed in this cell type (https://www.gtexportal.org/home/gene/KCTD11) and was therefore undetectable. To circumvent this, we generated plasmids encoding either wild-type (CMV-Myc-lKCTD11-WT) or mutant (CMV-Myc-lKCTD11-p.(Leu165Phefs9), CMV-Myc-lKCTD11-p.(Arg201Alafs50), and CMV-Myc-lKCTD11-p.(Arg73Trp)) Myc-tagged versions of lKCTD1, In HEK293 cells overexpressing these lKCTD11 proteins, we observed by Western blot, significantly reduced levels of all mutant lKCTD11 proteins as compared to the WT: a 58% decrease for Myc-lKCTD11p.(Leu165Phefs*9), an 81% decrease for Myc-lKCTD11p.(Arg201Alafs*50), and a 64% decrease for Myc-lKCTD11p.(Arg73Trp) (Fig. 2B and C). This result cannot be attributed to differences in transfection efficiency (Supplementary Fig.1A and B). We then investigated whether this reduction was due to post-transcriptional or post-translational mechanisms. Given that KCTD11 is a single exon gene and therefore lacks the exon-exon junctions required for Nonsense-Mediated mRNA Decay (NMD) in mammals, this mechanism is unlikely to be involved in the here (Cusack, Arndt et al. 2011, Karousis and Muhlemann 2019, Carrard and Lejeune 2023). Indeed, using semi-quantitative RT-PCR in HEK293 cells overexpressing WT or mutant lKCTD11 new observed no significant differences in transcript levels between WT and mutant constructs under identical transfection conditions(Supplementary Fig.1C), confirming that the observed decrease levels in mutant lKCTD11 protein result from post-translational rather than post-transcriptional mechanisms. Therefore, we investigated the contribution of autophagy and proteasome degradation to protein clearance by treating transfected HEK293 cells with either bafilomycin A1, an autophagy inhibitor, or MG-132, a proteasome inhibitor. Autophagy inhibition significantly increased mutant protein levels compared to untreated conditions, with a 78% increase for the p.(Leu165Phefs*9) mutant (2.8 ± 0.8), an 88% increase for the p.(Arg201Alafs*50) mutant (3.4 ± 1.2), and 81% for the p.Arg73Trp mutant (3 ± 1; Fig. 2D and E). In contrast, proteasome inhibition with MG-132 did not result in any significant increase in mutant protein levels compared to untreated conditions (Supplementary Fig.1D and E). Altogether, these results demonstrate the variants identified in KCTD11 are pathogenic and induce a loss of function, through enhanced degradation of the mutant proteins by autophagy

**Figure 2.**
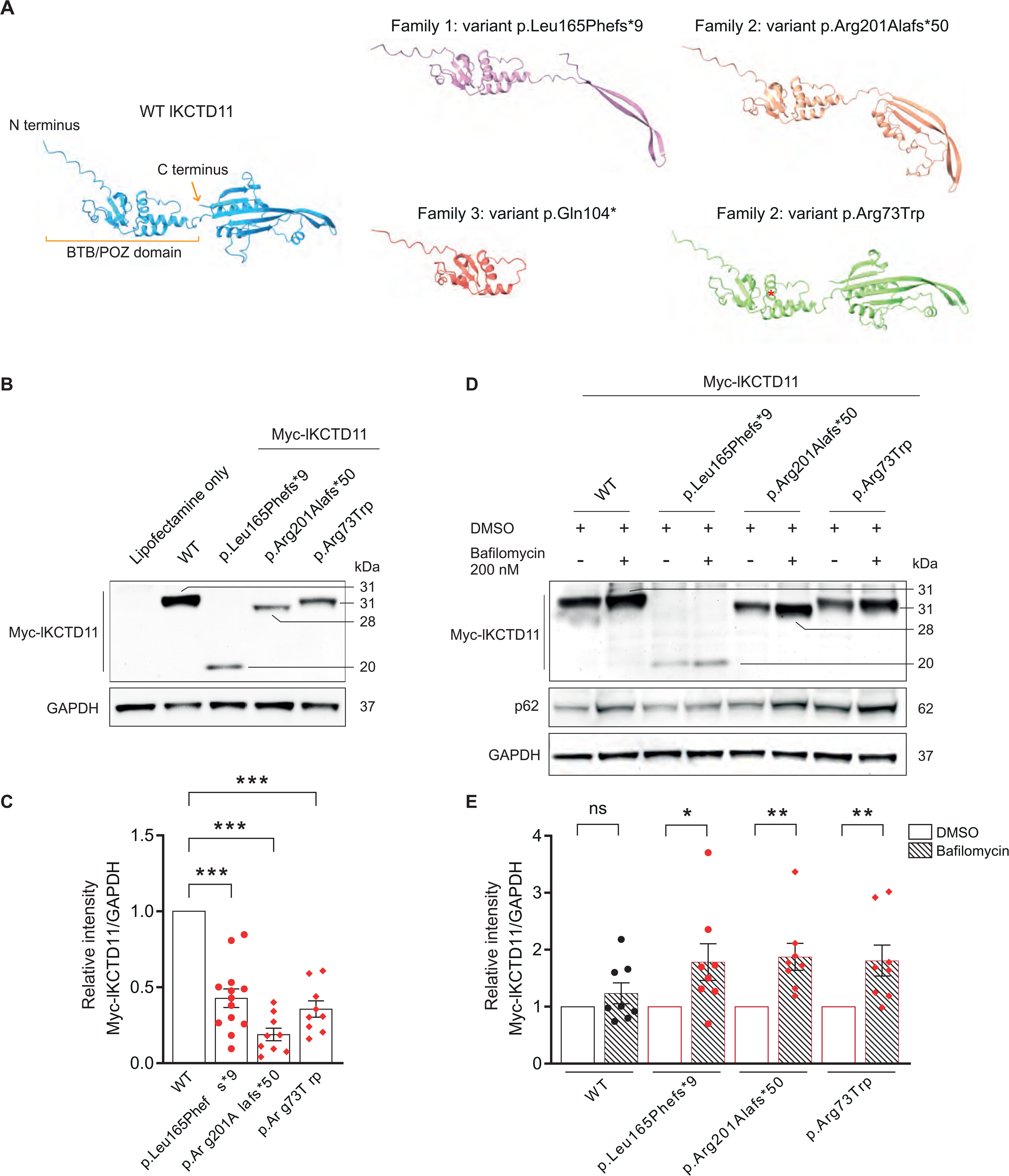
KCTD11 mutations are loss-of-function, inducing the post-translational degradation of the protein. (**A**) Three-dimensional structural model of KCTD11 long isoform (lKCTD11) protein (WT and mutant forms) generated using the AlphaFold2 AI system. Structures illustrate the predicted impact of identified mutations on KCTD11 folding. (**B**) Western blot analysis of Myc-tagged WT and mutant lKCTD11 proteins expressed in HEK293 cells, detected using an anti-Myc antibody. The p.(Leu165Phefs*9) mutation induces the production of a truncated protein of 99 amino acids, detected at around 20 kDa with the Myc-tag. The p.(Arg201Alafs*50) mutation results in a truncation of 22 amino acids, detected at approximately 28 kDa counting the Myc-tag. The p.(Arg73Trp) mutation does not significantly impact molecular weight, the protein being detected at 31 kDa counting the Myc-tag. (**C**) Quantification of relative intensity of Myc-tagged lKCTD11 proteins from western blot analysis in B, normalized against GAPDH. Data are expressed as mean ± SEM (n = 13 for p.(Leu165Phefs*9) mutation, n=9 for p.(Arg201Alafs*50) and p.(Arg73Trp) mutations; independent cultures) and normalized to the control (wild-type) value. Statistical analysis: one-way ANOVA, with Sidak *post hoc* test (multiple comparison to WT control). (**D**) Western blot analysis form HEK293 cells expressing the indicated construct (as in **A)** treated with vehicle control (DMSO) or 200 nM Bafilomycin A1 (autophagy inhibitor) for 6 hours. Autophagy inhibition efficacy was confirmed by assessing accumulation of p62 (SQSTM1). (**E**) Quantification of western blot results shown in (**D**). Data are expressed as mean ± SEM (n = 8 independent culture). Statistical analysis: unpaired Student’s *t*-test. ^∗^P < 0.05, ^∗∗^P < 0.01 and ^∗∗∗^P < 0.001.

### *KCTD11* is expressed in the peripheral nervous system

KCTD11 is a widely expressed protein, but is notably abundant in the PNS (https://www.gtexportal.org/home/gene/KCTD11). However, its role in the PNS remains largely unknown. We investigated its expression patterns as an initial step toward understanding its function.

We first investigated the expression profile of *Kctd11* during peripheral nerve development in mouse sciatic nerves using bulk mRNA sequencing at multiple postnatal stages: P0, P5, P15, and P30. Our findings ((expressed as transcripts per million, TPM) revealed a progressive increase in *Kctd11* expression throughout nerve development, reaching peak levels at P30, which corresponds to the young adult stage when developmental myelination is complete (Fig. 3A). Similarly, using TPM (Transcripts per million) normalized counts from Bulk mRNA sequencing in WT mouse DRGN/SC co-cultures at different time points, we show that the transcript levels of *kctd11* show a strong increase after the induction of myelination at Day 6, during active phases of myelination, an stabilization at later stages corresponding to myelin maintenance (mean TPM of 21.1 at Day 6, 41.3 at Day 14, and 43.5 at Day 22) (Fig. 3B).

**Figure 3.**
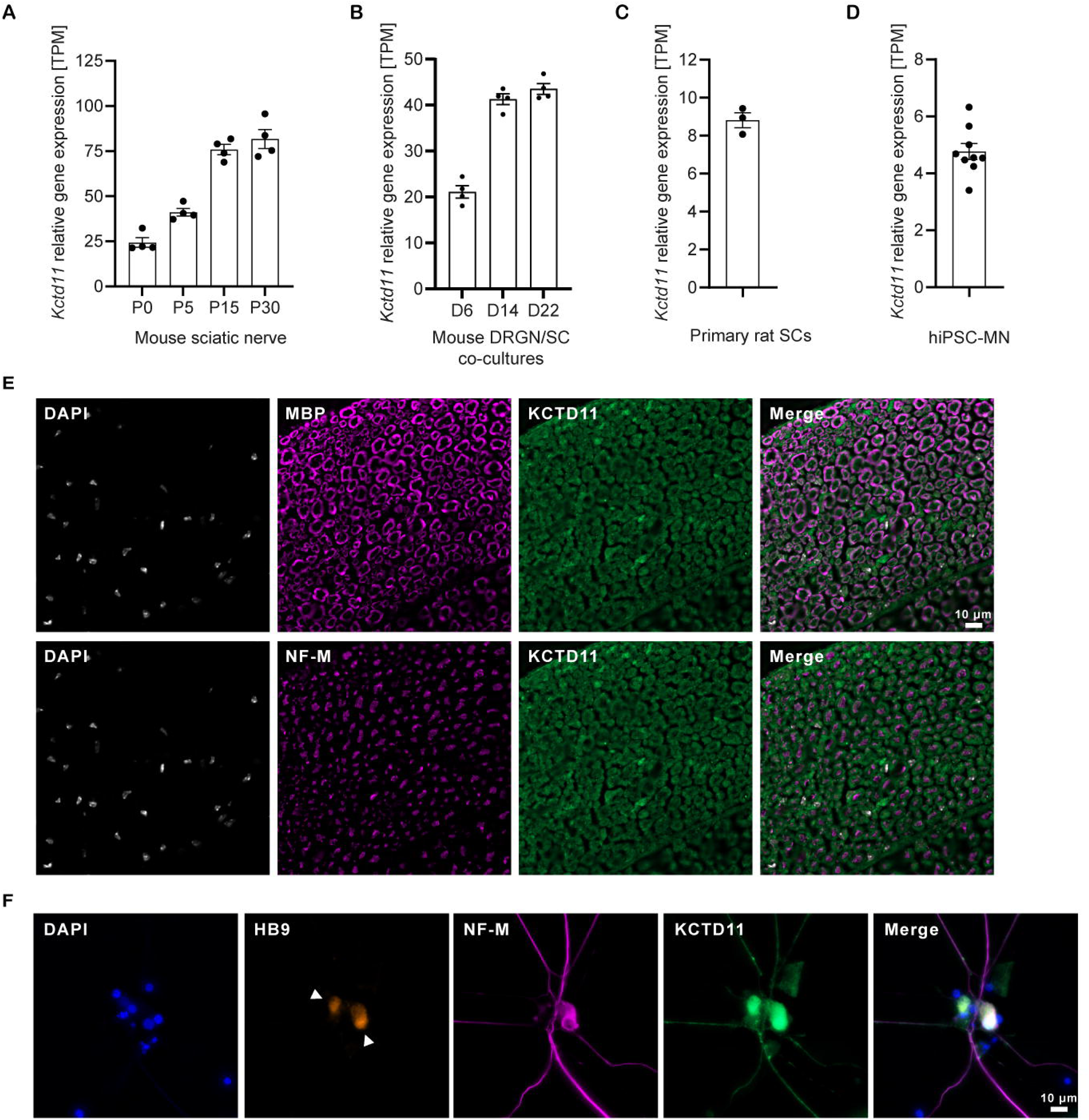
KCTD11 expression in the peripheral nervous system. (**A**) Kctd11 expression levels across peripheral nerve developmental stages, measured as Transcripts Per Million (TPM) using bulk mRNA sequencing data form WT mouse sciatic nerves at P0, P5, P15, and P30 (n = 4 per time point). (**B**) *Kctd11* expression levels in WT DRGN/SC co-culture samples at Day 6, Day 14, and Day 22, measured in Transcripts Per Million (TPM) from bulk mRNA sequencing data (n = 4 per time point). (**C**) Kctd11 expression levels in rat primary Schwann cells were measured as TPM using bulk mRNA sequencing data (n = 3). (**D**) Kctd11 expression levels in human induced pluripotent stem cell-derived motor neurons (hiPSC-MN) were measured as TPM using bulk mRNA sequencing data of eight controls. (**E**) Immunohistochemical of KCTD11 in cross-sections of sciatic nerve from WT mice, co-stained with NF-M (Neurofilament-M; neuronal projection marker) and MBP (Myelin Basic Protein, myelin marker). (**F**) Immunofluorescence analysis of KCTD11 in control hiPSC-derived mature motor neurons, co-stained with NF-M, and HB9 (Motor neuron marker).

Since the sciatic nerve is primarily composed of Schwann cells and axons, we tried to assess the relative contribution of each compartment by analysing bulk mRNA-Seq data from immature, non-myelinating, rat primary Schwann cells, and human induced pluripotent stem cell-derived motor neurons (hiPSC-derived MNs). Although we know that comparison of TPM values across samples might be biased, the levels of *kctd11* transcripts seem higher in SCs than in Motor Neurons (mean TPM values= 8.8 for primary rat Schwann cells and 4.2 for hiPSC-MNs,Fig. 3C-D).

Immunostaining of KCTD11 in sciatic nerve cross-sections from 3-month-old (P90) mice (Fig. 3D), showing expression in both the axon and the myelin sheath, with higher expression of KCTD11 in the latter, as evidenced by its colocalization with Myelin Basic Protein (MBP), and a weaker signal detected in axons, labeled by Neurofilament medium (NF-M) staining (Fig. 3E). In the neurons, KCTD11 is primarily localized to the soma, but is also detected in neurites as demonstrated by colocalization with HB9, a motor neuron-specific nuclear marker, and NF-M respectively, in human induced pluripotent stem cell-derived motor neurons (hiPSC-MN) (Fig. 3F). Collectively, these findings show that, in the PNS, KCTD11 is expressed both in neurons and in the myelin,, particularly at stages where myelin is mature and maintenance processes are established,

### Loss of KCTD11 disrupts myelination dynamics *in vitro*

In order to further investigate how the loss of KCTD11 impacts PNS function, we used and *in vitro* myelin model derived from our *Kctd11^-/ -^*knock-out mouse model: it is based on primary cell co-culture of sensory dorsal root ganglion (DRG) neurons, with Schwann cells (SC), derived from E13.5 embryos DRGs of *Kctd11^-/-^* or WT mice .

We first confirmed the knockout by performing bulk mRNA sequencing on sciatic nerves from *Kctd11^-/-^* and WT mice (Supplementary Fig. 3A). Due to the lack of effective antibodies, we were unfortunately unable to validate the knockout at the protein level.

We next investigated the impact of *kctd11* ablation on PNS myelination in DRGN/SC co-cultures from E13.5 *Kctd11^-/-^* and WT embryos. We first evaluated the impact of KCTD11 loss on axonal morphology of DRG neurons alone, by removing Schwann cells, after dissection of the DRG: immunofluorescent labeling of neurites revealed no noticeable morphological differences between *Kctd11^-/-^* and WT DRG neurons (Fig.4A). Then, we evaluated myelination processes/dynamics in the DRGN/SC cocultures, through quantification of MBP positive myelin segments, representing myelinating SCs, at various end time points (D14, D18 and D22: respectively 8, 12 and 16 days after the induction of myelinationby acid ascorbic addition at day 6 of coculture). We did not observe any morphological myelin abnormalities in *Kctd11^-/-^* co-cultures as compared to WT, at any stage (Fig. 4B). However, quantitative analysis revealed significantly fewer myelin segments in *Kctd11^-/-^* DRGN/SC cocultures as compared to control at Day 14 (*Kctd11^-/-^*: 126 ± 20 vs. WT: 196 ± 47; 44% decrease) and Day 18 (*Kctd11^-/-^*: 304 ± 29 vs. WT: 316 ± 79; 42% decrease) (Fig.4C and D). Interestingly, this pattern was reversed at Day 22, with *Kctd11^-/-^*DRGN/SC cocultures displaying a significantly higher number of myelin segments as compared to WT (*Kctd11^-/-^*: 2479 ± 440 vs. WT: 2441 ± 267; 112% increase) (Fig.4E). At Day 14, myelin segments were also shorter in *Kctd11^-/-^* DRGN/SC cocultures as compared to the WT (Fig.4F).

**Figure 4.**
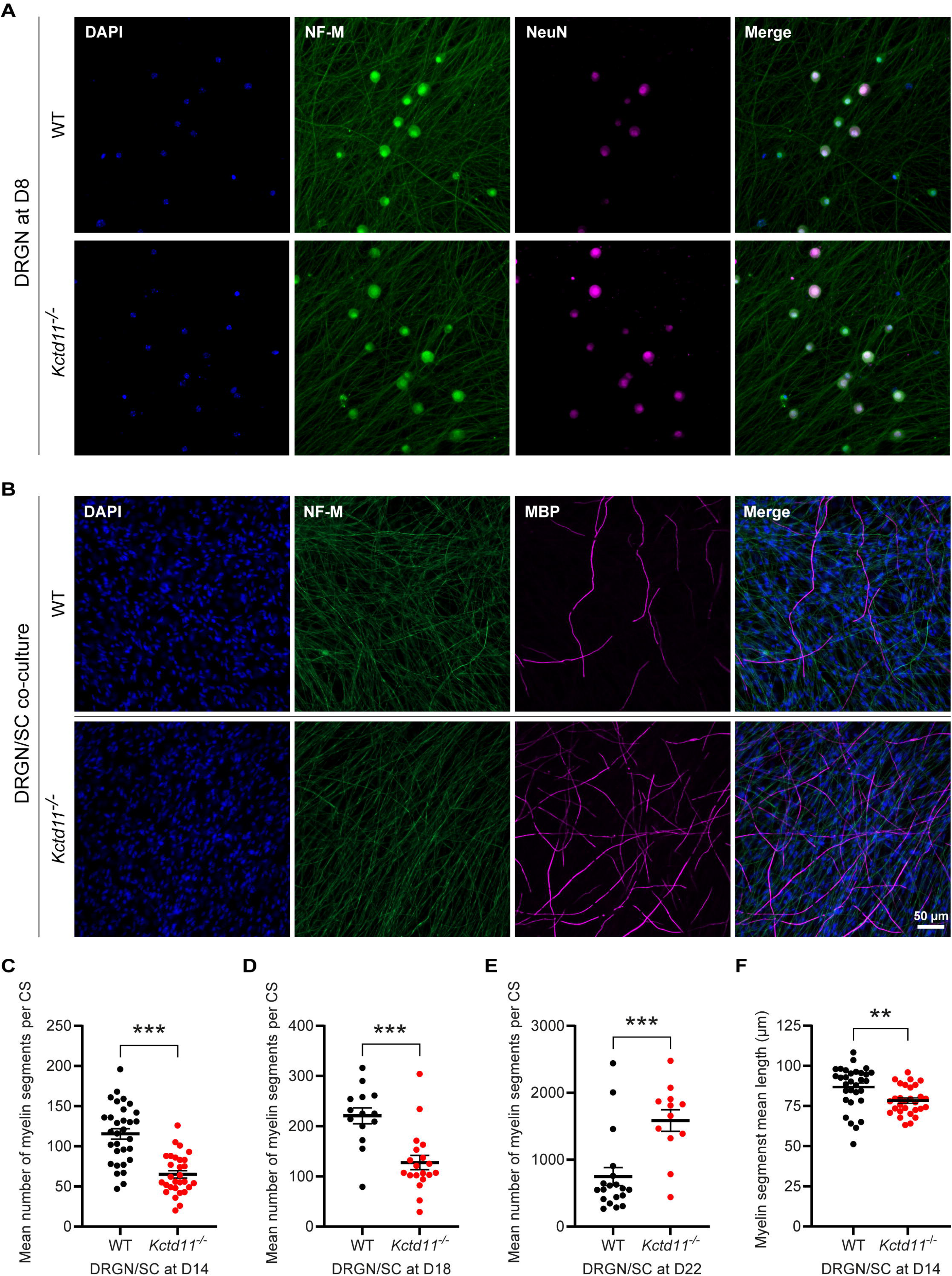
The loss of KCTD11 leads to impaired myelination *in vitro*. (**A**) Immunostaining of neurites (NF-M) in cultured DRG neurons (DRGN) derived from *Kctd11^−/−^* and WT mice, co-stained with NeuN (neuronal marker). (**B**) Representative image of myelinated fibers 16 days after induction of myelination (D22) in *Kctd11^−/−^* and WT DRGN/SC co-cultures. Myelinating Schwann cells were identified by MBP immunostaining, while DRGN neuronal fibers were labeled with an NF-M antibody. (**C-E**) Quantification of myelin segment number in *Kctd11*^−/−^ and WT DRGN/SC co-cultures at (**C**) D14, (**D**) D18, and (**E**) D22. Myelin segments were identified by MBP immunostaining. CS: coverslip. Statistical analysis was performed using two-way ANOVA with Sidak’s post hoc test (multiple comparisons: mean WT versus *Kctd11*^−/−^ at each time point). (**F**) Quantification of myelin segment length (µm) in WT and *Kctd11*^−/−^ DRGN/SC co-cultures at D14. A minimum of 30 segments per coverslip were measured. Statistical analysis was performed using an unpaired Student’s *t*-test. ^∗^P < 0.05, ^∗∗^P < 0.01 and ^∗∗∗^P < 0.001.

### Loss of KCTD11 disrupts PNS myelination *in vivo*

In parallel, we explored the impact of the loss of KCTD11 in the PNS physiology *in vivo*, through histological, electrophysiological, and behavioral characterization of the *Kctd11^-/-^* mouse model versus WT mice . Given the progressive nature of CMT disease, we carried out our study over time in *Kctd11^-/-^* and control mice, aged 3, 6, 12 and 18 months. At the histopathological level, we studied the distal part of several nerves: the sciatic nerve (a proximal mixed sensory and motor nerve), the tibial nerve (a more distal mixed sensory and motor nerve), and the saphenous nerve (very distal, pure sensory nerve) by electron microscopy (Fig.5).

**Figure 5.**
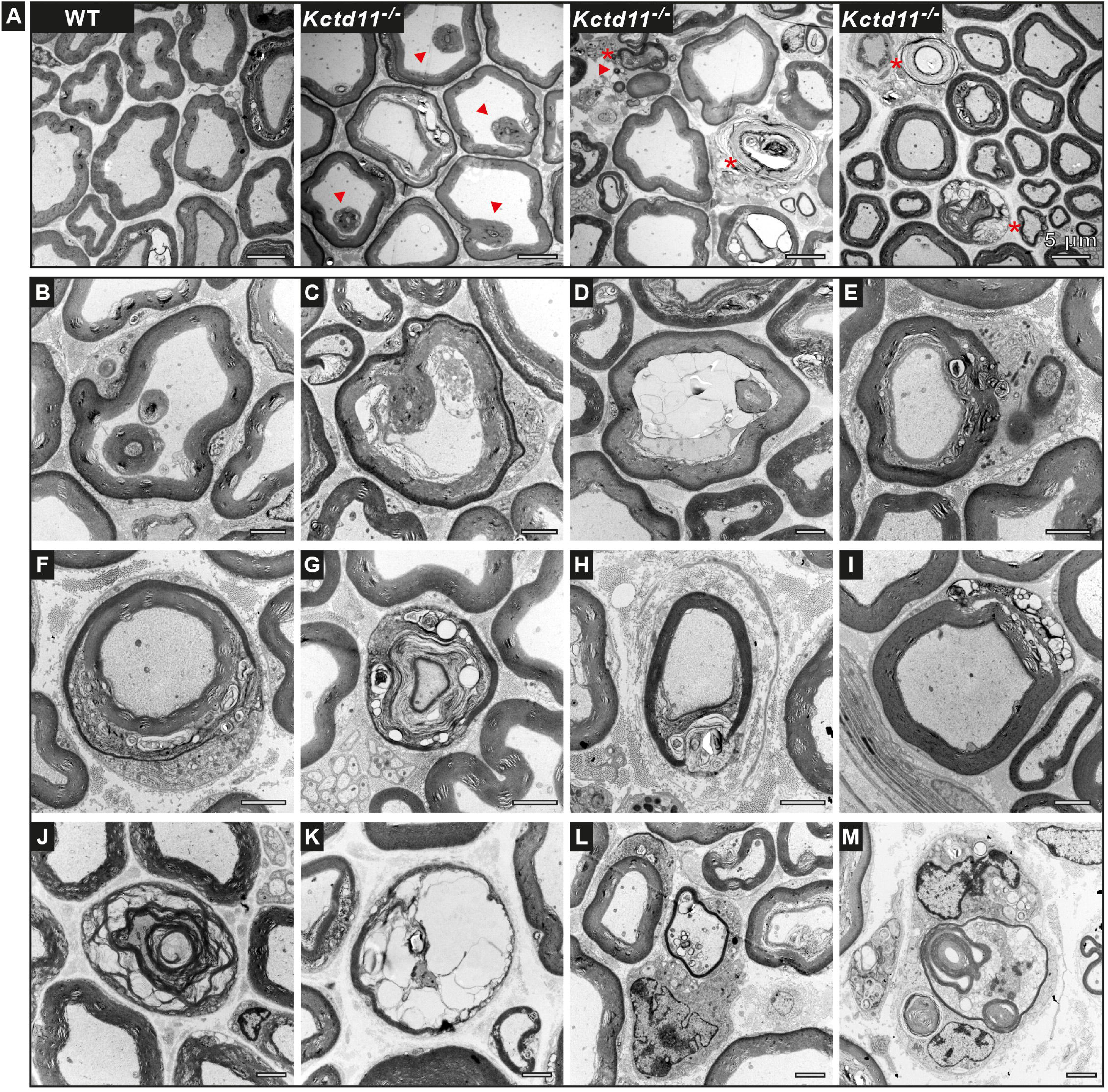
KCTD11 loss induce impaired myelination *in vivo*: *Kctd11^-/-^* mice display abnormal myelin features at 18 months. (**A**) Electron microscopy images of cross-sections from the distal region of the sciatic nerve in 18-month-old WT and *Kctd11^-/-^* mice. Arrowheads indicate myelin abnormalities. Asterisks indicate myelin segments showing signs of degradation. **(B–M)** Electron microscopy images of sciatic nerve cross-sections from 18-month-old *Kctd11^-/-^* mice. (**B–E**) Abnormally myelinated fibers exhibiting early signs of myelin degradation: (**B**) myelin infolding; (**C**) myelin infolding with multivesicular disintegration of adaxonal myelin lamellae; (**D**) infolding accompanied by large vacuoles within Schwann cell adaxonal myelin lamellae; (**E**) myelin outfolding with vacuolization at the level of the Schmidt-Lanterman incisure. (**F–I**) Fibers displaying early signs of myelin degradation without overt myelin abnormalities. (**J, K)** Fibers with advanced myelin degradation, and **(L, M**) fibers with advanced myelin degradation accompanied by remyelination. Scale: 2 µM.

We observed abnormal myelin structures, mostly infoldings, revealing aberrant myelination, in all *Kctd11^-/-^* nerves at different ages (Fig.6). The proportion of those abnormalities was higher in the sciatic and tibial nerves of *Kctd11^-/-^* mice as compared to WT, as early as 3 months old, with a significant increase over time (in the sciatic nerve: 3-month-old: WT 6.3 ± 2.5 % and *Kctd11^-/-^* 10.3 ± 8.2 %; 6-month-old: WT 10 ± 5.4 % and *Kctd11^-/-^* 15.5 ± 11 %; 12-month-old: WT 8.2 ± 3.5 % and *Kctd11^-/-^* 16.7 ± 8.8 %; 18-month-old: WT 7.3 ± 4.2 % and *Kctd11^-/-^* 17.1 ± 9.6 %; in the tibial nerve: 3-month-old: WT 8 ± 4.6 % and *Kctd11^-/-^* 10.2 ± 7.8 %; 6-month-old: WT 10.4 ± 5.2 % and *Kctd11^-/-^* 16.2 ± 10.1 %; 12-month-old: WT 7.7 ± 2.2 % and *Kctd11^-/-^* 14.6 ± 8.5 %; 18-month-old: WT 8.1 ± 5 % and *Kctd11^-/-^* 17.3 ± 12.1%) (Fig.6A and B). In the saphenous nerve, the proportion of abnormally myelinated fibers was significantly higher only in older *Kctd11^-/^* mice (12- and 18-month-old)*^-^* (3-month-old: WT 12 ± 4.7 % and *Kctd11^-/-^* 15.7 ± 10.9 %; 6-month-old: WT 11.3 ± 9.3 % and *Kctd11^-/-^* 15.5 ± 12.4 %; 12-month-old: WT 8.8 ± 5 % and *Kctd11^-/-^*23.9 ± 10.1 %; 18-month-old: WT 8 ± 4.6 % and *Kctd11^-/-^* 17.8 ± 12 %) (Fig. 6C).

**Figure 6.**
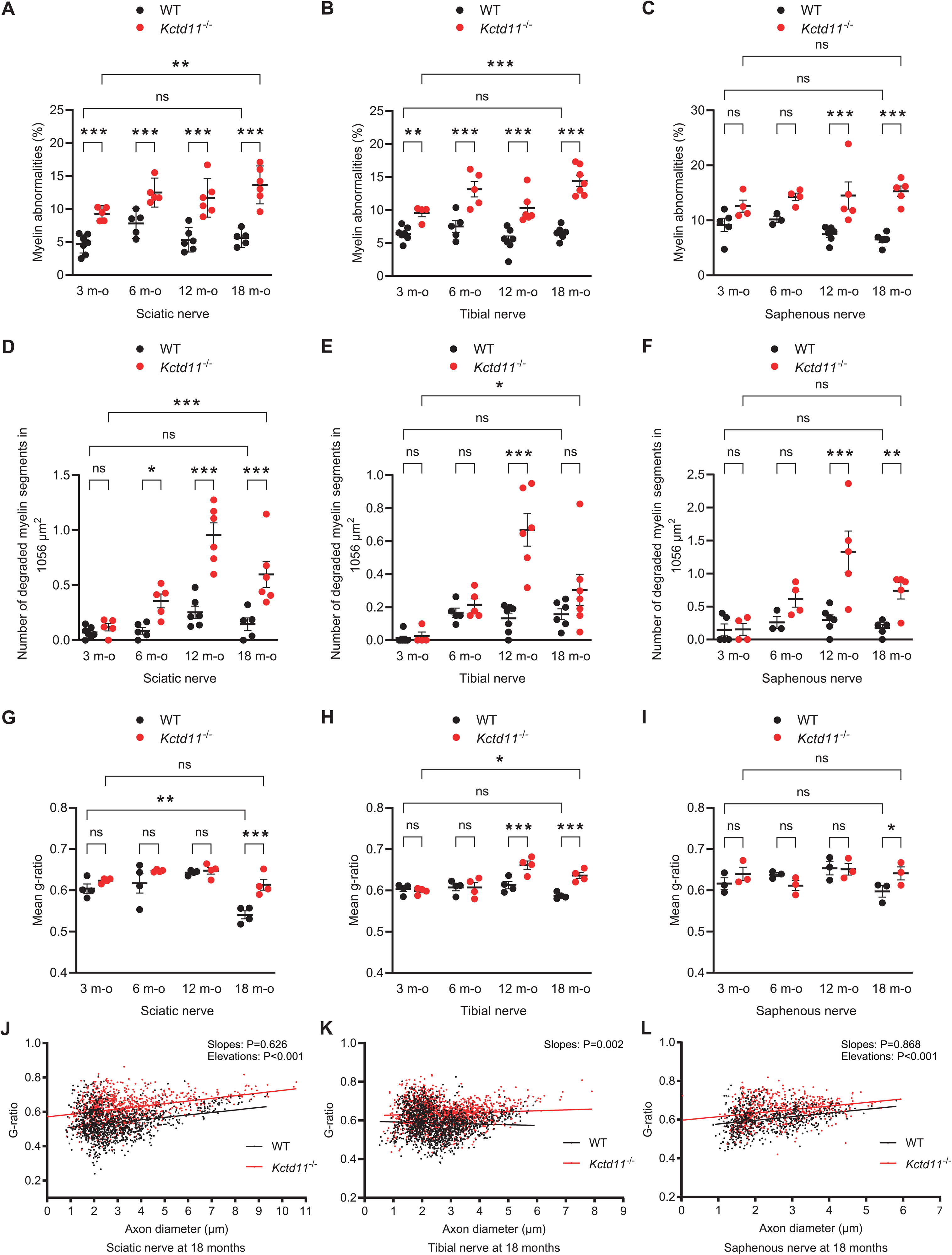
Histological characterization of a novel *Kctd11^-/-^* mouse model of intermediate CMT disease. (**A-C**) Quantification of the proportion of abnormally myelinated fibers in the distal regions of: (**A**) the sciatic nerve (containing both motor and sensory fibers), (**B**) the tibial nerve (more distal, containing both motor and sensory fibers), and (**C**) the saphenous nerve (purely sensory), from 3-, 6-, 12-, and 18-month-old *Kctd11*^−/−^ and WT mice (*n* = 4–7 animals per genotype). Data are expressed as mean percentage ± SEM. Statistical analysis: two-way ANOVA with Tukey’s post hoc test for multiple comparisons. Myelin abnormalities are defined as myelin infoldings. (**D-F**) Quantification of myelin segments showing degradation within a 1056 µm² nerve area in distal regions of the (**D**) sciatic, (**E**) tibial, and (**F**) saphenous nerves of 3-, 6-, 12-, and 18-month-old *Kctd11*^−/−^ and WT mice (*n* = 4–7 animals per genotype). Data are expressed as mean percentage ± SEM. Statistical analysis: two-way ANOVA with Tukey’s post hoc test for multiple comparisons. (**G**-**I**) G-ratio analyses of nerve fibers from (**G**) sciatic, (**H**) tibial, and (**I**) saphenous nerves of 3-, 6-, 12-, and 18-month-old *Kctd11*^−/−^ and WT mice (sciatic and tibial nerves: *n* = 4 per genotype; saphenous nerve: *n* = 3 per genotype). A higher g-ratio indicates thinner myelin. A total of 1,000 to 2,000 axons, ranging from 0.5 to 9 µm in diameter, were analyzed per genotype and per age group. Data are presented as mean ± SEM. Statistical analysis: two-way ANOVA with Tukey’s post hoc test for multiple comparisons. (**J-L**) Scatter plots representing the g-ratio (y-axis) of individual axons as a function of their respective diameters (x-axis) in the distal regions of the (**J**) sciatic, (**K**) tibial, and (**L**) saphenous nerves from 18-month-old *Kctd11*^−/−^ and WT mice. Each point corresponds to an individual fiber. ∗P < 0.05, ∗∗P < 0.01, and ∗∗∗P < 0.001.

Additionally to the presence of aberrant myelin structures, we evidenced increased myelin degradation in the mutant nerves. Indeed, the number of degraded myelin segments was higher in *Kctd11^-/-^*sciatic nerves as compared to controls, as soon as 6-month-old, and increased significantly over time (3-month-old: WT 0.15 ± 0 and *Kctd11^-/-^* 0.19 ± 0; 6-month-old: WT 0.17 ± 0 and *Kctd11^-/-^* 0.52 ± 0.17; 12-month-old: WT 0.48 ± 0.13 and *Kctd11^-/-^* 1.28 ± 0.6; 18-month-old: WT 0.32 ± 0 and *Kctd11^-/-^*1.15 ± 0.35) (Fig. 6D). In the tibial and saphenous nerves, we also observed a significant increase of degraded myelin segments, but they were significant starting from 12-month-old (in the tibial nerve: 3-month-old: WT 0.08 ± 0 and *Kctd11^-/-^* 0.1 ± 0; 6-month-old: WT 0.26 ± 0.15 and *Kctd11^-/-^* 0.33 ± 0.15; 12-month-old: WT 0.21 ± 0 and *Kctd11^-/-^* 0.95 ± 0.32; 18-month-old: WT 0.25 ± 0.1 and *Kctd11^-/-^* 0.83 ± 0.05; in the saphenous nerve: 3-month-old: WT 0.64 ± 0.59 and *Kctd11^-/-^* 0.67 ± 0.62; 6-month-old: WT 0.65 ± 0.63 and *Kctd11^-/-^* 0.63 ± 0.59; 12-month-old 0.68 ± 0.62 and *Kctd11^-/-^* 0.68 ± 0.63; 18-month-old: WT 0.61 ± 0.57 and *Kctd11^-/-^* 0.67 ± 0.61; Fig. 6D-F).

Myelin thickness was assessed by g-ratio (axon diameter/fibre diameter) measurements on myelinated axons without overt myelin aberrations in the three studied nerves, at all time points. No significant changes were noticed before age 18 months, where, in all nerves, a significant increase of the mean g-ratio was observed in mutant mice as compared to WT was observed at older age (18 months old) (in the sciatic nerve: 3-month-old: WT 0.63 ± 0.58 and *Kctd11^-/-^* 0.63 ± 0.62; 6-month-old: WT 0.66 ± 0.55 and *Kctd11^-/-^* 0.65 ± 0.64; 12-month-old: WT 0.65 ± 0.63 and *Kctd11^-/-^* 0.66 ± 0.62; 18-month-old: WT 0.56 ± 0.52 and *Kctd11^-/-^* 0.65 ± 0.59; in the tibial nerve: 3-month-old: WT 0.61 ± 0.58 and *Kctd11^-/-^* 0.60 ± 0.59; 6-month-old: WT 0.62 ± 0.58 and *Kctd11^-/-^* 0.63 ± 0.58; 12-month-old: WT 0.64 ± 0.60 and *Kctd11^-/-^* 0.68 ± 0.63; 18-month-old: WT 0.60 ± 0.58 and *Kctd11^-/-^* 0.65 ± 0.62; in the saphenous nerve: 3-month-old: WT 0.64 ± 0.59 and *Kctd11^-/-^* 0.67 ± 0.62; 6-month-old: WT 0.65 ± 0.63 and *Kctd11^-/-^* 0.63 ± 0.59; 12-month-old 0.68 ± 0.62 and *Kctd11^-/-^* 0.68 ± 0.63; 18-month-old: WT 0.61 ± 0.57 and *Kctd11^-/-^* 0.67 ± 0.61; Fig.6G-I and 6J-L).. This alteration was not correlated to differences in the axonal diameter (Supplementary Fig. 5A-C).

However, to also clarify the role of KCTD11 during development, we studied the effect of its ablation on sciatic nerve development using immunohistological and teased fiber analyses (data not shown), which revealed no significant effects, suggesting that KCTD11 may primarily function during later stages associated with peripheral nerve maintenance.

To assess if the loss of KCTD11 induces any measurable behavioral abnormalities in the knock-out mice, we first reviewed the results published by the International Mouse Phenotyping Consortium (IMPC, https://www.mousephenotype.org). Indeed, the knock-out mouse model that we use, has been generated in the frame of the KOMP program (Birling, Yoshiki et al. 2021) and therefore has been extensively characterized through this IMPC Consortium (Groza, Gomez et al. 2023). In their phenotyping characterization, there was no striking phenotype, except increased erythrocyte cell number and haemoglobin content. In particular they showed no difference between the mutant homozygous *Kctd11^-/-^*mice and the control mice, in the Grip Strength test, the only test assessing defects of the neuromuscular system, nor in the open field test which assesses locomotor activity globally (https://www.mousephenotype.org/data/genes/MGI:2448712). However, there are several limitations to IMPC study: i) although the control group is very large; the mutant group comprises only 8 mice (4 males/4 femelles); ii) the studied mice were early adults and there is no longitudinal follow-up. Therefore, to further assess the locomotor performance of our knock-out mice, we performed a longitudinal study using 3 tests: Rotarod to test motor function, Grip strength test to assess muscle strength and Gait analysis for analysis of motor coordination and synchrony (Brooks and Dunnett 2009), in 3 groups of age an sex matched animals (n>14 animals in each group) at age 6-12 and 18 months. We also measured the Nerve Conduction Velocities, longitudinally, in knock-out (*Kctd11^-/-^*, n= 5) versus Controls (WT n= 5) mice, at age 12 and 18 months. Our results confirm the results obtained by the IMPC and show no behavioral or electrophysiological deficits in *Kctd11^-/-^* mice as compared to heterozygous or WT controls (Supplementary Fig. 6D-F). These results are not surprising for models of Charcot-Marie-Tooth disease, especially considering the moderate phenotype in the patients.

### Loss of KCTD11 leads to deregulation of HDAC1 and the Wnt/β-catenin pathway *in vitro* and *in vivo*

To explore the pathophysiological mechanisms underlying the defects observed *in vitro* in the absence of KCTD11, we investigated known partners of KCTD11, involved in myelination and the balance between cell proliferation and differentiation, specifically HDAC1, β-catenin, and components of the mTORC1 pathway (mTOR, S6, p-S6, 4EBP1, p-4EBP1)(Jacob, Christen et al. 2011, Figlia, Gerber et al. 2018, Maurice and Angers 2025). In vitro, we detected no apparent differences in the expression or localization of HDAC1, β-catenin, or mTORC1 pathway proteins in primary cultures of DRG neurons (DRGN) lacking KCTD11 as compared to control DRG neurons (data not shown).

We next focused our investigation on DRGN/SC co-cultures at two distinct culture stages (days 14 and 22: 8 and 16 days after the induction of myelination respectively),. At day 14, there were no significant differences for any of the tested proteins (Fig.7 and Supplementary Fig.4). In contrast, at day 22, while we detected no differences in the protein and phosphorylation levels of proteins from the mTORC1 pathway (Supplementary Fig.4), we observed a statistically significant decrease in β-catenin phosphorylation accompanied by reduced HDAC1 levels (Fig.7A-D). This finding suggests a late perturbation of the Wnt/β-catenin signaling pathway, potentially linked to altered myelin maintenance, when KCTD11 is absent. In vivo, in sciatic nerves from old 18 month-old mice, we observed the opposite tendency, i.e. a significant increase in the protein levels of total β-catenin and HDAC1, with no change in the levels of phosphorylated β-catenin (Fig.7E-G). No changes were detected in the mTOR and AKT pathways (Supp Fig4H-I)

**Figure 7.**
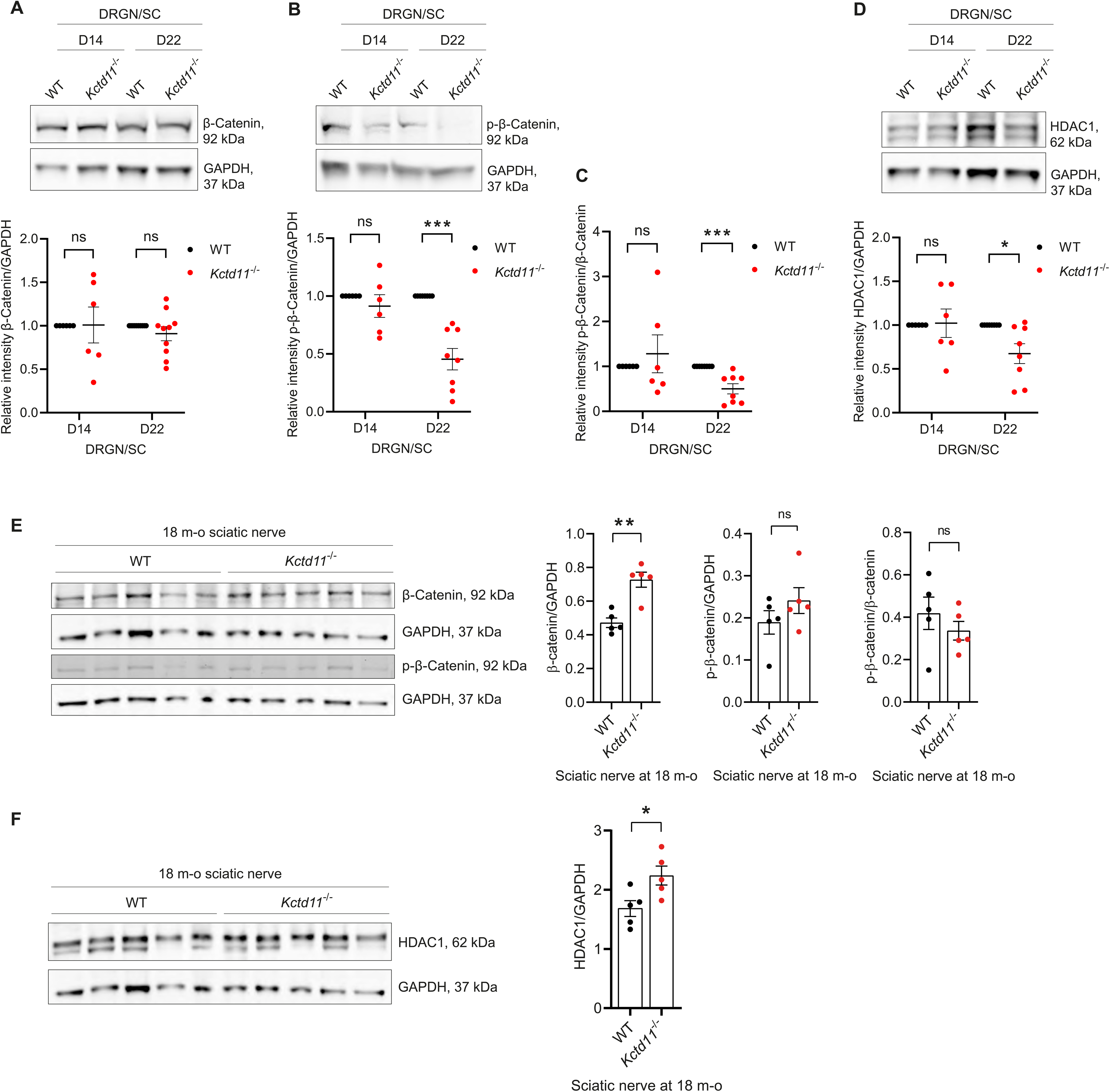
Wnt/β-Catenin pathway is dysregulated in *Kctd11^-/-^* DRGN/SC co-cultures (A-D) and sciatic nerves (E-F). **(A)** Top: Representative Western blot images showing β-Catenin expression in *Kctd11^−/−^* and WT DRGN/SC co-cultures at D14 and D22. Bottom: Quantification of β-Catenin expression levels normalized to GAPDH. Data are expressed as mean ± SEM (D14: n = 6; D22: n = 10). Statistical analysis: unpaired Student’s *t*-test. **(B)** Top: Representative Western blot images showing phosphorylated β-Catenin (p-β-Catenin) expression in *Kctd11^−/−^* and WT DRGN/SC co-cultures at D14 and D22. Bottom: Quantification of p-β-Catenin relative intensity normalized to GAPDH. Data are expressed as mean ± SEM (D14: n = 6; D22: n = 9). Statistical analysis: unpaired Student’s *t*-test. **(C)** Quantification of the ratio of phosphorylated to total β-Catenin (p-β-Catenin/β-Catenin) normalized to GAPDH in *Kctd11^−/−^* and WT DRGN/SC co-cultures at D14 and D22. Data are expressed as mean ± SEM (D14: n = 6; D22: n = 8). Statistical analysis: unpaired Student’s *t*-test. (**D**) Top: Representative Western blot images showing HDAC1 expression in *Kctd11^−/−^*and WT DRGN/SC co-cultures at D14 and D22. Bottom: Quantification of HDAC1 relative intensity normalized to GAPDH. Data are expressed as mean ± SEM (D14: n = 6; D22: n = 8). Statistical analysis: unpaired Student’s *t*-test. P < 0.05, ^∗∗^P < 0.01 and ^∗∗∗^P < 0.001. **E)** Left: Representative Western blot images showing β-Catenin and phosphorylated β-Catenin (p-β-Catenin) expression in sciatic nerves from 18 months-old *Kctd11^−/−^* and WT mice . Right: Quantification of β-Catenin and p-β-Catenin expression levels normalized to GAPDH as well as normalized p-β-Catenin/β-Catenin ratio. Data are expressed as mean ± SEM (n=5 for each genotype). Statistical analysis: unpaired Student’s *t*-test. **F)** Left: Representative Western blot images showing HDAC1 expression in sciatic nerves from 18 months-old *Kctd11^−/−^* and WT mice . Right: of HDAC1 relative intensity normalized to GAPDH. Data are expressed as mean ± SEM (n=5 for each genotype). Statistical analysis: unpaired Student’s *t*-test.

Overall, these data suggest an increased Wnt/ β-catenin signaling pathway.

### Loss of KCTD11 alters the transcriptional profile and dynamic in *in vitro* myelin samples

We showed *in vitro* and *in vivo* that the absence of KCTD11 leads to altered myelination and myelination dynamics. To further decipher the mechanisms involved in these alterations, we assessed global gene expression changes resulting from the loss of KCTD11, by performing bulk mRNA sequencing on DRGN/SC co-cultures from *Kctd11^-/-^* and WT samples, at three different time points: (i) culture day 6 (D6), representing the initiation stage before the induction of myelination with ascorbic acid; (ii) culture day 14 (D14), an intermediate stage after 8 days of myelination induction (14 days of total co-culture); and (iii) culture day 22 (D22), a late stage following 16 days of myelination induction.

Differential Gene Expression analysis, using an adjusted P-value< 0.05, revealed very few differentially expressed genes (DEGs) at Day 6 and D14 (Supplementary Fig.6A),with only five deregulated genes at Day 6 (three upregulated and two downregulated), and 62 at Day 14 (27 upregulated and 35 downregulated) (Fig.8A). However, at Day 22, a total of 4,322 DEGs were identified (adjusted P-value< 0.05) (Fig. 8A-B), strengthening the hypothesis of a role for KCTD11 in later stages of myelination and myelin maintenance *in vitro.* Among the deregulated genes, 1846 were upregulated and 2476 downregulated (Fige. 8A).

**Figure 8.**
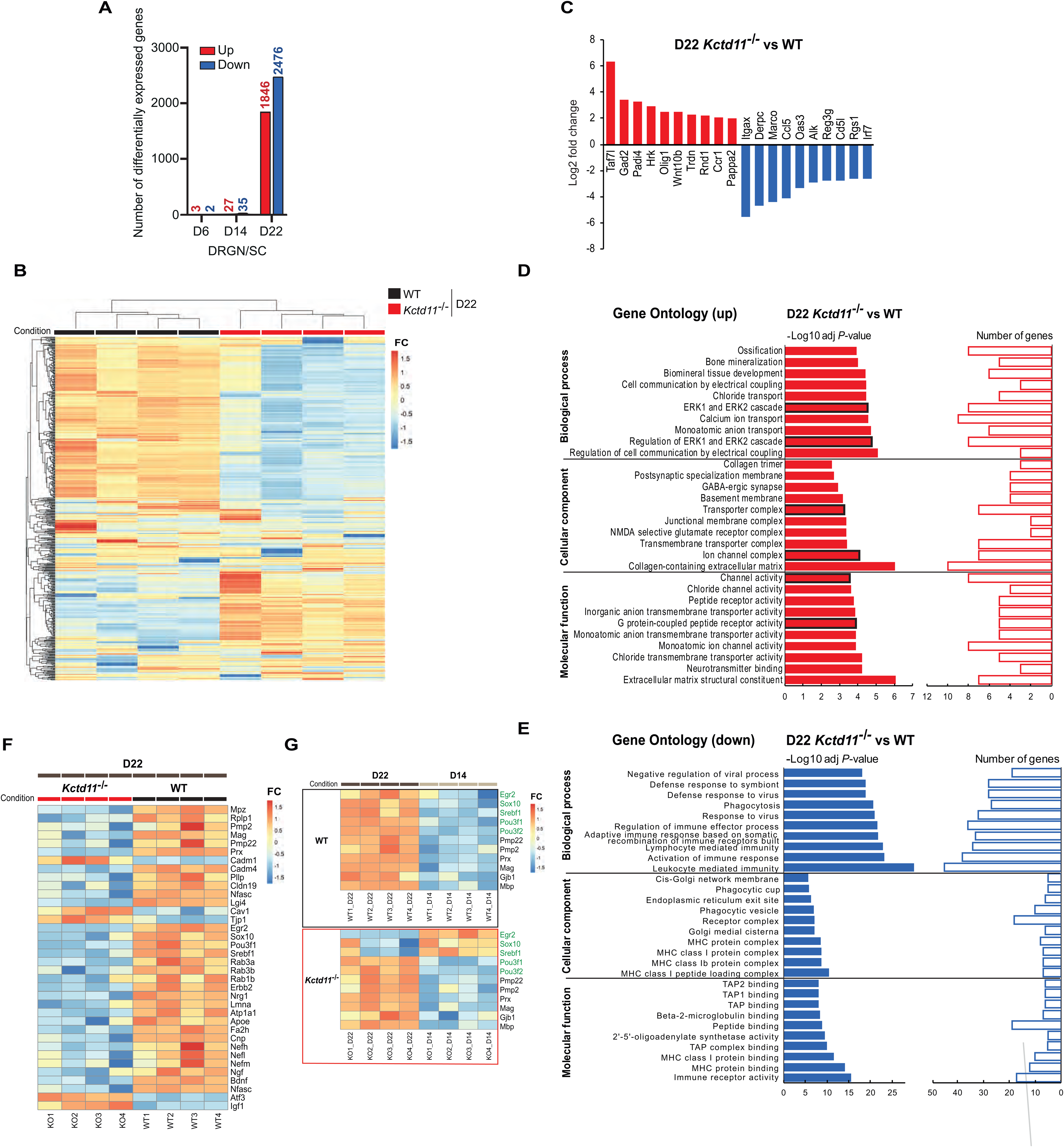
Bulk transcriptomic (mRNA-Seq) analysis of *in vitro* myelin samples (DRGN/SC co-cultures) from knock-out (*Kctd11*^−/−^) and control (WT) mice. (**A**) Number of differentially expressed genes (DEGs) identified by Bulk mRNA sequencing in knock-out versus control DRGN/SC co-cultures (n=4 replicates for *kctd11^-/-^* and 4 replicates for WT) at each time point, with adjusted p-value < 0.05 and fold change (FC) > 2 (upregulated) or < 0.5 (downregulated). (**B**) Full heat map of unsupervised hierarchical clustering of the eight samples at Day 22 (n = 4 replicates for *Kctd11^-/-^* and n = 4 replicates for WT). The scale bar represents values obtained after applying a variance-stabilizing transformation to the count data (DESeq2: VarianceStabilizingTransformation) before normalization. High expression is indicated in red, while low expression is shown in blue. (**C**) Top 10 upregulated (showed in red) and downregulated (showed in blue) genes with an adjusted P-value < 0.01 identified by Differential Gene Expression analysis in knock-out versus control DRGN/SC cocultures at Day 22 cultures (n=4 replicates for *kctd11^-/-^* and 4 replicates for WT). (**D-E**).Gene Ontology (GO) enrichment analysis of Differentially Expressed Genes (DEGs) in knock-out versus control DRGN/SC cocultures at Day 22 cultures (n=4 replicates for *kctd11^-/-^*and 4 replicates for WT). *Left bars represent the enrichment score (–Log10 of adjusted P-value) for each GO term. GO terms with particularly relevant functions related to the myelination process or/and neuronal function are outlined in black. Right bars represent the number of genes counted for each GO term* (**D**) Top 10 Biological Process (BP), Cellular Component (CC), and Molecular Function (MF) categories identified from **the upregulated transcripts** (DEGs with an adjusted *P*-value < 0.01 and a FC > 2). (**E**) Top 10 BP, CC and MF categories identified from **the downregulated transcripts** (DEGs with an adjusted *P*-value < 0.01 and a FC < 0.5. Left bars represent the enrichment score (–Log10 of adjusted *P*-value) for each GO term, while right bars indicate the number of genes associated for each GO term. (**F**) Heatmap illustrating Differential Gene Expression (adjusted P-value < 0.05) of myelin-related genes in *Kctd11^-/-^* (n = 4 replicates) versus WT (n = 4 replicates) DRGN/SC coculture samples at D22. Expression intensity is indicated by color (warm colors: high expression, cool colors: low expression). (**G**) Heatmaps illustrating Differential Gene Expression (adjusted P-value < 0.05) of myelin-related genes at DRGN/SC coculture Day 22 versus DRGN/SC coculture D14 (n = 4 replicates at each time point) either D14 or D22 in *Kctd11^-/-^* or WT samples. Key transcription factors driving Schwann cell differentiation are listed in green, while major structural myelin proteins of Schwann cells are listed in black. Expression intensity is indicated by color (warm colors: high expression, cool colors: low expression).

In the *Kctd11*^−/−^ versus WT co-cultures at Day 22, among the top 10 upregulated or downregulated genes (*Padj* value < 0.01, FC>2 or FC < 0.5) (Fig. 8C), we found several members of the Wnt/β-catenin pathway, such as *Wnt10a* (upregulated, FC=5.51, padj= 0.00283) and *Alk* (downregulated, FC=0.134, padj= 7.21×10^−19^); genes encoding proteins involved in cell proliferation/apoptosis (*Hrk*, *Derpc*, *Reg3g* and *Padi4*). Interestingly, *Padi4*, a Peptidylarginine deiminase, converting a methylarginine residue to citrulline at specific sites on the tails of histones H3 and H4, has been shown to collaborate with HDAC1 to generate a repressive chromatin environment on promoters (Denis, Deplus et al. 2009).

Functional Gene Ontology (GO) annotation of the DEGs found at Day 22, with statistical significance of Padj value<0.01, FC>2 (upregulated) or <0.5 (downregulated), showed enrichment in several Cellular Component (CC), Biological Process (BP), and Molecular Function (MF). For upregulated genes, in the top10 enriched BP GO terms, w e found ‘ERK1 and ERK2 cascade’ (GO: 0070371), ‘Regulation of ERK1 and ERK2 cascade’ (GO: 0070372)(Fig. 8D), which are related to Schwann cells myelination. We also identified GO terms related to neuronal function, such as ‘Ion channel complex’ (GO: 0034702), ‘Channel activity’ (GO: 0015267), and ‘Monatomic ion channel activity’ (GO: 0005216) (Fig. 8C). Functional GO annotation of the downregulated DEGs, with a significance threshold of Padj < 0.01 and fold change < 0.05, revealed enrichment in pathways related to immune functions, MHC complex and TAP Binding (Fig.8E). This can be explained by the fact that, to initiate myelin clearance by macrophages, Schwann cells reportedly have the capability to act as facultative antigen-presenting cells, expressing MHC complex II molecules (Bergsteinsdottir et al., 1992; Meyer zu Hörste et al., 2010).

Considering the strong and significant increase in the number of myelin segments in the *Kctd11*^−/−^ cocultures at Day 22 as compared to controls, we compared the expression levels of the major myelination-related genes in the RNA-Seq dataset at Day 22. As expected,. The heatmap of differentially expressed genes (D22 KO vs. WT), filtered for known myelination-related genes (FC < 0.8 or > 1.2, Padj < 0.05), indicates a strong downregulation for most genes, and upregulation only for the following genes: Cav1, Atf3, Tjp1, Igf1, Cam1), probably as an attempt to stop enhanced myelination in DRGN/SC cocultures in the Knock-out mice (Fig.8E). Comparison of the expression of those genes between Day 22 and Day 14 in each genotype shows that the dynamics of the induction of expression of the major transcriptional regulators of Schwann cell myelination (*Egr2, Sox10, Srebf1*) is altered in the KO DRGN/SC cocultures (Fig.8F).

### Loss of KCTD11 shows deregulations of the Wnt/β-catenin, Sonic Hedgehog and Hippo/YAP pathways

mRNA-Seq transcriptomic analyses were also performed, in quadriplicate, on total RNA extracted from tibial nerves of *Kctd11^-/-^* or WT mice at age 12 and 18 months. Few genes were deregulated (padj<0.05) either at 12 months (356 up, 253 down) or at 18 months ((155 up, 23 down) (Supplementary Figure 6B), therefore functional analysis of the DEGs choosing a FC threshold of 2 for upregulation and 0.5 for downregulation did not allow a functional analysis of these data. However, detailed analysis of the DEGs in 12 month-old mice, showed very interesting deregulations in tibial nerves from *Kctd11^-/-^* mice as compared to controls. Among upregulated transcripts (FC>2, padj value<0.05), we found *Gli2* (pAdj ∼ 1,0 × 10^-12^ ; FC ∼ 2,47), involved in the sonic Hedgehog (shh) pathway, negatively regulated by KCTD11; *Wnt10a* (pAdj ∼ 0,039 ; FC ∼ 2,41), a member of the Wnt/ β-catenin pathway, also negatively regulated by KCTD11, *Ccn2* (*CTGF*) (padj ∼ 4,04 × 10^-10^ ; FC ∼ 3), target of YAP/TAZ, which expression is induced when the Hippo/YAP pathway is inhibited, which is the case in the absence of KCTD11 here. At the opposite, we noticed a downregulation of *Cxxc4* (padj ∼ 0.038; FC ∼ 0,39), an inhibitor of the Wnt/β-catenin pathway.

In nerves from 18-month old of *Kctd11^-/-^* mice, the most deregulated transcript is *Igkc* (FC ∼ 489, padj∼0.014). Together with *C6* (FC∼6.89, padj∼0.0134) and *S100a8* (FC∼6.58, padj∼0.0046), they are implicated in immune surveillance and inflammation.

## Discussion

Charcot-Marie-Tooth disease is a heterogenous group of diseases of the Peripheral Nervous System, particularly at the genetic level, with more than 100 causative genes identified to date. Based on the measure of median motor nerve conduction velocity (MNCV), we distinguish demyelinating CMT1 (with MNCV <38 m/s), axonal CMT2 (MNCV >38 m/s), and intermediate CMT with MNCV between 25 and 45 m/s (Berciano, Garcia et al. 2017). In this study, we identify *KCTD11* as a novel causative gene for autosomal recessive Intermediate CMT (RI-CMT), thereby expanding the genetic landscape of inherited neuropathies and bringing to five the number of genes involved in RI-CMT. We first identified a homozygous c.493del p.(Leu165Phefs*9) pathogenic frameshift variant in *KCTD11* in six patients from a large consanguineous Palestinian family from Lebanon. Subsequent identification of 4 additional patients from 4 unrelated families with diverse ethnic background (French, Italian and Egyptian), with biallelic variants in *KCTD11*, confirmed its implication as a new CMT gene. All patients share the same clinical presentation of rather late-onset, slowly progressive, moderate, predominantly motor, intermediate form of Charcot-Marie-Tooth disease. In younger patients, the MNCVs at the median nerve are close to 45 m/s and might be interpreted as axonal, but based on the definition by Berciano et al. (MNCVs between 25 and 45 m/s at the median nerve), we describe this new form of autosomal recessive CMT as intermediate RI-CMTE.

We demonstrate, *in vitro*, that the identified variants are pathogenic and induce loss of function through enhanced degradation of the mutated KCTD11 proteins by autophagy. KCTD11, also named REN, was first described as a developmentally regulated gene that promotes neural cell differentiation (Gallo, Zazzeroni et al. 2002). Known as a tumor suppressor gene (Di Marcotullio, Ferretti et al. 2004, Zazzeroni, Nicosia et al. 2014, Tong, Yang et al. 2017), its role in the PNS myelination had never been explored before. In order to better understand how the loss of KCTD11 function in the patients lead to the peripheral neuropathy, we sought to investigate the role of KCTD11 in the PNS using both *in vitro* and *in vivo* models knock-out for the gene. Considering that the patients present an intermediate form of CMT, the involvement of both Schwann cells and peripheral neurons is likely.

*In vivo*, we studied knock-out *Kctd11^-/-^* mice from birth to the age 18 months. As usually observed for mouse models of Charcot-Marie-Tooth disease (Bosco, Falzone et al. 2021), we observed no overt phenotype despite extensive longitudinal behavioral studies from age 6 to 18 months. However, at the histopathological level, the knock-out mice exhibit abnormal myelin structures, mostly infoldings, as soon as 3 months of age, which increase over age, as well as signs of myelin degradation at older ages, particularly in the distal regions of the sciatic, tibial, and saphenous nerves. Very interestingly, mice deficient for *Plekhg5*, another RI-CMT gene (RI-CMTC, MIM 615376) (Azzedine, Zavadakova et al. 2013), which is also responsible for dHMN (MIM 611067) (Maystadt, Rezsohazy et al. 2007), show defective axon/Schwann cell units characterized by myelin infoldings in peripheral nerves (Luningschror, Slotta et al. 2020). This excessive myelin production usually precedes demyelination and axon loss and therefore, the histopathological defects observed in our *Kctd11^-/-^* mice are in accordance with the fact that our patients are affected with intermediate/axonal CMT. The significant but moderate number of anomalies, increasing over age, and the observation that mixed nerves (sciatic and tibial) were more affected than the purely sensory saphenous nerve, are also in line with the predominant motor impairment, and a middle/late disease onset in the patients. The similarity in the nerve abnormalities between *kctd11* and *plekhg5* deficient mice might be representative of common pathophysiological mechanisms. Given the role of PLEKHG5 in regulating autophagy (Luningschror, Binotti et al. 2017), the fact that *Kctd11* deficient mice nerves have increased myelin degradation, therefore possible altered myelinophagy (Gomez-Sanchez, Carty et al. 2015) and our observation that the mutant KCTD11 proteins seem to be degraded by autophagy, it is tempting to speculate that there might be a link between KCTD11 and PLEKHG5 and therefore common pathomechanisms between those two CMT diseases. This hypothesis remain to be investigated both *in vivo* and *in vitro*.

*In vitro*, in DRG neurons, the ablation of KCTD11 did not induce any detectable cellular defects. However, in the myelin model derived from the knock-out *Kctd11^-/-^* mouse (DRGN/SC cocultures), we confirmed defects in the myelination process, more precisely in myelination dynamics, although we did not observe the myelin abnormalities seen *in vivo*. Myelination seems to be slowed or delayed during the first days of myelination, while, at the opposite, it gets out of control, leading to increased myelination, in later stages corresponding to myelin maintenance. Data extracted from mRNA-seq on these DRGN/SC cocultures confirm altered temporal modulation of the expression of the major transcriptional myelination regulators EGR2 and SOX10 between Day 14 and Day 22 in *Kctd11^-/-^* mice as compared to controls. However, some time points (Day 18) are missing to fully explain the temporal deregulations and will be explored in the future.

Mechanistically, both at the RNA and the protein level, *in vitro* and *in vivo*, our result point to activation of Wnt/β-catenin signaling, detected by decreased ratio of phosphorylated β-catenin/total β-catenin (statistically significant *in vitro*, tendency *in vivo*), either by lowering the levels of phosphorylated inactive β-catenin as seen *in vivo*, or by increasing the levels of total β-catenin as observed *in vivo*. Unfortunately, our antibodies against β-catenin were not specific to active β-catenin (ABC), and therefore, we can only deduce from our results that we have an increase in ABC levels due to the loss of KCTD11. This might affect the balance between proliferation, differentiation and apoptosis in Schwann cells with loss of function mutations in KCTD11.Our results suggest a role in myelin maintenance for KCTD11, rather than in earlier steps of myelin development as well as the initiation of a repair transcriptional program similar to the one observed during neuroinflammation or nerve injury.

Additionally, very interestingly, *in vitro* RNA-Seq suggest also activation of the Sonic Hedgehog signaling pathway and inhibition of the Hippo/YAP. Those pathway are closely related and mutually regulate each other and are also regulated by HDAC1, by limiting the levels of ABC (Jacob, Christen et al. 2011).

The precise mechanisms by which the loss of KCTD11 induces this modulation of these pathways is yet unclear, and further experiments are needed *in vitro* and *in vivo*, in particular in order to assess the role of autophagy (Petherick, Williams et al. 2013, Lorzadeh, Kohan et al. 2021) in the pathomechanisms involved. The discrepancies between the *in vitro* and *in vivo* models might be due to the fact that the observed time points are not exactly similar, whereas the observed processes are very tightly regulated over time.

Overall, we showed that our *Kctd11^-/-^* mouse and its derived in vitro myelin model, are reliable tools to study the molecular mechanisms underlying aberrant myelination in RI-CMTE. However, considering the likely involvement of both Schwann cells and the peripheral neurons in the disease pathophysiological mechanisms, we are aware that the used complete knock-out model does allow to decipher the contribution of each PNS component individually. Therefore, we will need to build conditional SC, motor neuron or DRG neuron knock-out models to further study the specific role of KCTD11 in each part of the PNS. As an alternative, in vivo, we intend to study specifically the axon component in motor neurons or DRG neurons derived from patients’ hiPSC, at our disposal as previously done for other CMT subtypes (El-Bazzal, Rihan et al. 2019, Bos, Rihan et al. 2022).

In conclusion, our findings establish KCTD11 as a novel gene responsible for RI-CMTE. We show that KCTD11 is a key regulator of myelination, via Schwann cell dysfunction and possible axonal defects.Our results highlight the role of defective Wnt/β-catenin, Hippo/YAP and Shh signaling pathways, at least in Schwann cells. Future studies are needed to precisely decipher the molecular mechanisms, however our results bring new insights into the genetics and pathophysiology of Charcot-Marie-Tooth disease by describing KCTD11 as a new player in PNS myelination.

## Acknowledgements

We thank the PiCSL-FBI core facility (IBDM, AMU-Marseille), member of the France BioImaging national research infrastructure, in particular Aïcha Aouane and Nicolas Brouilly, who performed the electron microscopy experiments. We are grateful to Catherine Aubert from the Genomics and Bioinformatics facility (GBiM), from the U 1251/Marseille Medical Genetics laboratory, for supervising the bulk mRNA-seq experiments. We are grateful to Mahdi Ghozal for his technical help in the overexpression experiments. This research was made possible through access to the data generated by the France Genomic Medicine Plan 2025. Finally, we would like to thank Mazen Bou Malhab for his help in desiging the figures.

## Supplementary material

Supplementary material is available at *Brain* online.

## Appendix 1

This section is included if the article contains an appendix, for example, to list consortium collaborators who must be indexed online.

## Legend to supplementary Figures and Tables

**Supplementary figure 1**

(**A**) Quantification of the proportion of Myc-positive HEK293 cells transfected with either Myc-lKCTD11c.493del (p.(Leu165Phefs*9)) or Myc-lKCTD11-WT construct (n=13 independent cultures). Data are expressed as mean percentage ± SEM. Statistical analysis: unpaired Student’s *t*-test.

(**B**) Quantification of the proportion of Myc-positive HEK293 cells transfected with either Myc-lKCTD11-c.601del (p.(Arg201Alafs*50)), Myc-lKCTD11-c.217C>T (p.(Arg73Trp)) or Myc-lKCTD11-WT constructs (n=9 independent cultures). Data are expressed as mean percentage ± SEM. Statistical analysis: one-way ANOVA, with Sidak *post hoc* test (multiple comparison to WT control).

(**C**) RT-PCR amplification of *Myc-tagged lKCTD11* and *ACTB* (NM_001101) transcripts from HEK293 cells transfected with indicated constructs. The expected amplicon size is 800 bp for both *Kctd11* mouse isoforms and 246 bp for *ACTB* transcripts.

(**D**) Western blot analysis form HEK293 cells expressing the indicated construct treated with vehicle control (DMSO) or 5 µM MG-132 (proteasome inhibitor) for 6 hours. Proteasome inhibition efficacy was confirmed by assessing accumulation of ubiquitinated protein.

(**E**) Quantification of western blot results shown in (**D**). Data are expressed as mean ± SEM (n ≥ 5 independent culture). Statistical analysis: unpaired Student’s *t*-test. ^∗^P < 0.05, ^∗∗^P < 0.01 and ^∗∗∗^P < 0.001.

**Supplementary figure 2**

(**A**) Immunohistochemical analysis of KCTD11 in cross-sections of spinal cord from WT mice, co-stained with NeuN (neuronal marker). Abbreviation: VH: ventral horn; DH: dorsal horn.

**Supplementary figure 3**

(**A**) Genomic deletion of KCTD11 in the *Kctd11^-/-^*mouse model confirmed by IGV visualization of BAM files from bulk mRNA-sequencing of sciatic nerves from *Kctd11^-/-^* and WT mice. In WT samples, reads fully cover the *Kctd11* coding sequence (100 % coverage; sequencing depth: 50×). In contrast, no reads were detected in the *Kctd11^-/-^* sample, confirming the absence of Kctd11 transcripts and validating the efficiency of the *Kctd11*-knockout in our mouse model. Data aligned on Mouse Genome Reference: GRCm39/mm39.

**Supplementary figure 4**

(**A**) Top: Representative Western blot images showing mTOR expression in *Kctd11^−/−^* and WT DRGN/SC co-cultures at D14 and D22. Bottom: Quantification of mTOR relative intensity normalized to GAPDH. Data are expressed as mean ± SEM (D14: n = 6; D22: n = 11). Statistical analysis: unpaired Student’s *t*-test.

(**B**) Top: Representative Western blot images showing S6 expression in *Kctd11^−/−^* and WT DRGN/SC co-cultures at D14 and D22. Bottom: Quantification of S6 relative intensity normalized to GAPDH. Data are expressed as mean ± SEM (D14: n = 6; D22: n = 8). Statistical analysis: unpaired Student’s *t*-test.

(**C**) Top: Representative Western blot images showing p-S6 expression in *Kctd11^−/−^* and WT DRGN/SC co-cultures at D14 and D22. Bottom: Quantification of p-S6 relative intensity normalized to GAPDH. Data are expressed as mean ± SEM (D14: n = 6; D22: n = 11). Statistical analysis: unpaired Student’s *t*-test.

(**D**) Quantification of the ratio of phosphorylated to total S6 (p-S6/S6) normalized to GAPDH in *Kctd11^−/−^*and WT DRGN/SC co-cultures at D14 and D22. Data are expressed as mean ± SEM (D14: n = 6; D22: n = 6). Statistical analysis: unpaired Student’s *t*-test.

(**E**) Top: Representative Western blot images showing 4EBP1 expression in *Kctd11^−/−^* and WT DRGN/SC co-cultures at D14 and D22. Bottom: Quantification of 4EBP1 relative intensity normalized to GAPDH. Data are expressed as mean ± SEM (D14: n = 6; D22: n = 9). Statistical analysis: unpaired Student’s *t*-test.

(**F**) Top: Representative Western blot images showing p-4EBP1 expression in *Kctd11^−/−^* and WT DRGN/SC co-cultures at D14 and D22. Bottom: Quantification of p-4EBP1 relative intensity normalized to GAPDH. Data are expressed as mean ± SEM (D14: n = 6; D22: n = 10). Statistical analysis: unpaired Student’s *t*-test.

(**G**) Quantification of the ratio of phosphorylated to total 4EBP1 (p-4EBP1/4EBP1) normalized to GAPDH in *Kctd11^−/−^* and WT DRGN/SC co-cultures at D14 and D22. Data are expressed as mean ± SEM (D14: n = 6; D22: n = 8). Statistical analysis: unpaired Student’s *t*-test. P < 0.05, ^∗∗^P < 0.01 and ^∗∗∗^P < 0.001.

(**H**) Left: Representative Western blot images showing mTOR and p-4EBP1expression in sciatic nerves from 18 months-old *Kctd11^−/−^* and WT mice. Right: Quantification of mTOR and p-4EBP1 relative intensity normalized to GAPDH. Data are expressed as mean ± SEM (n = 5 in each gentype). Statistical analysis: unpaired Student’s *t*-test.

(**I**) Left: Representative Western blot images showing Akt and p-Akt expression in sciatic nerves from 18 months-old *Kctd11^−/−^*and WT mice. Right: Quantification of Akt and p-Akt relative intensity normalized to GAPDH, as well as the pAkt/Akt ratio. Data are expressed as mean ± SEM (n = 5 in each gentype). Statistical analysis: unpaired Student’s *t*-test.

**Supplementary figure 5**

(**A-C**) Quantification of the mean axon diameter (µm) in (**A**) sciatic, (**B**) tibial, and (C) saphenous nerves of *Kctd11*^−/−^ and WT mice at 3, 6, 12, and 18 months of age. A total of 1,000 to 2,000 axons per genotype and per age group, ranging from 0.5 to 9 µm in diameter, were analyzed. Data are presented as mean ± SEM. Statistical analysis was performed using two-way ANOVA with Tukey’s post hoc test (multiple comparisons). P < 0.05, ^∗∗^P < 0.01 and ^∗∗∗^P < 0.001.

**Supplementary figure 6 Transcriptional profiles of *in vitro* myelin samples (DRGN/SC co-cultures) or tibial nerves from knock-out (*Kctd11*^−/−^) and control (WT) mice over time**

**(A)** Full heat map of unsupervised hierarchical clustering of all 24 samples (n = 4 replicates for *Kctd11^-/-^* and n = 4 replicates for WT at each coculture Day 6, 14 and 22). The scale bar represents values obtained after applying a variance-stabilizing transformation to the count data (DESeq2: VarianceStabilizingTransformation) before normalization. High expression is indicated in red, while low expression is shown in blue

**(B)** Full heat map of unsupervised hierarchical clustering of all 16 samples (n = 4 replicates for *Kctd11^-/-^* and n = 4 replicates for WT at each age: 12 and 18 months). The scale bar represents values obtained after applying a variance-stabilizing transformation to the count data (DESeq2: VarianceStabilizingTransformation) before normalization. High expression is indicated in red, while low expression is shown in blue

**Supplementary figure 7**

**(A)** Left: Representative Western blot images showing ErbB2 expression in *Kctd11^−/−^* and WT DRGN/SC co-cultures at D22. Right: Quantification of ErbB2 expression levels normalized to GAPDH. Data are expressed as mean ± SEM (D22: n = 11). Statistical analysis: unpaired Student’s *t*-test.

**(B)** Left: Representative Western blot images showing phospho-ErbB2 expression in *Kctd11^−/−^* and WT DRGN/SC co-cultures at D22. Right: Quantification of phosphor-ErbB2 expression levels normalized to GAPDH. Data are expressed as mean ± SEM (D22: n = 7). Statistical analysis: unpaired Student’s *t*-test. P < 0.05, ^∗∗^P < 0.01 and ^∗∗∗^P < 0.001.

**Supplementary Table 1. Final list of homozygous by Descent Variants in Family 1**

Chr = Chromosome; Ref = Reference allele; Alt = alternate allele; Depth = number of reads at position; gnomAD_exome MAF = minor allele frequency in gnomAD exome dataset; gnomAD_genome MAF = minor allele frequency in gnomAD genome dataset; GME MAF = minor allele frequency in the GME dataset, N= Not relevant

**Supplementary Table 2 Sequencing statistics and filtering**

**A.** NGS sequencing coverage statistics. **B**. Details of filtering steps. **C.** MAFs of variants identified in KCTD11 (NM_001363642)

## References

Adzhubei, I. A., S. Schmidt, L. Peshkin, V. E. Ramensky, A. Gerasimova, P. Bork, A. S. Kondrashov and S. R. Sunyaev (2010). “A method and server for predicting damaging missense mutations.” Nature Methods 7(4): 248–249.

Alazami, A. M., N. Patel, H. E. Shamseldin, S. Anazi, M. S. Al-Dosari, F. Alzahrani, H. Hijazi, M. Alshammari, M. A. Aldahmesh, M. A. Salih, E. Faqeih, A. Alhashem, F. A. Bashiri, M. Al-Owain, A. Y. Kentab, S. Sogaty, S. Al Tala, M. H. Temsah, M. Tulbah, R. F. Aljelaify, S. A. Alshahwan, M. Z. Seidahmed, A. A. Alhadid, H. Aldhalaan, F. AlQallaf, W. Kurdi, M. Alfadhel, Z. Babay, M. Alsogheer, N. Kaya, Z. N. Al-Hassnan, G. M. Abdel-Salam, N. Al-Sannaa, F. Al Mutairi, H. Y. El Khashab, S. Bohlega, X. Jia, H. C. Nguyen, R. Hammami, N. Adly, J. Y. Mohamed, F. Abdulwahab, N. Ibrahim, E. A. Naim, B. Al-Younes, B. F. Meyer, M. Hashem, R. Shaheen, Y. Xiong, M. Abouelhoda, A. A. Aldeeri, D. M. Monies and F. S. Alkuraya (2015). “Accelerating novel candidate gene discovery in neurogenetic disorders via whole-exome sequencing of prescreened multiplex consanguineous families.” Cell Rep 10(2): 148–161.

Argenti, B., R. Gallo, L. Di Marcotullio, E. Ferretti, M. Napolitano, S. Canterini, E. De Smaele, A. Greco, M. T. Fiorenza, M. Maroder, I. Screpanti, E. Alesse and A. Gulino (2005). “Hedgehog antagonist REN(KCTD11) regulates proliferation and apoptosis of developing granule cell progenitors.” J Neurosci 25(36): 8338–8346.

Azzedine, H., P. Zavadakova, V. Plante-Bordeneuve, M. Vaz Pato, N. Pinto, L. Bartesaghi, J. Zenker, O. Poirot, N. Bernard-Marissal, E. Arnaud Gouttenoire, R. Cartoni, A. Title, G. Venturini, J. J. Medard, E. Makowski, L. Schols, K. G. Claeys, C. Stendel, A. Roos, J. Weis, O. Dubourg, J. Leal Loureiro, G. Stevanin, G. Said, A. Amato, J. Baraban, E. LeGuern, J. Senderek, C. Rivolta and R. Chrast (2013). “PLEKHG5 deficiency leads to an intermediate form of autosomal-recessive Charcot-Marie-Tooth disease.” Hum Mol Genet 22(20): 4224–4232.

Baranski, T. J., A. T. Kraja, J. L. Fink, M. Feitosa, P. A. Lenzini, I. B. Borecki, C. T. Liu, L. A. Cupples, K. E. North and M. A. Province (2018). “A high throughput, functional screen of human Body Mass Index GWAS loci using tissue-specific RNAi Drosophila melanogaster crosses.” PLoS Genet 14(4): e1007222.

Barreto, L. C., F. S. Oliveira, P. S. Nunes, I. M. de Franca Costa, C. A. Garcez, G. M. Goes, E. L. Neves, J. de Souza Siqueira Quintans and A. A. de Souza Araujo (2016). “Epidemiologic Study of Charcot-Marie-Tooth Disease: A Systematic Review.” Neuroepidemiology 46(3): 157–165.

Baux, D., C. Van Goethem, O. Ardouin, T. Guignard, A. Bergougnoux, M. Koenig and A. F. Roux (2021). “MobiDetails: online DNA variants interpretation.” Eur J Hum Genet 29(2): 356–360.

Berciano, J., A. Garcia, E. Gallardo, K. Peeters, A. L. Pelayo-Negro, S. Alvarez-Paradelo, J. Gazulla, M. Martinez-Tames, J. Infante and A. Jordanova (2017). “Intermediate Charcot-Marie-Tooth disease: an electrophysiological reappraisal and systematic review.” J Neurol 264(8): 1655–1677.

Berciano, J., A. Garcia, E. Gallardo, K. Peeters, A. L. Pelayo-Negro, S. Alvarez-Paradelo, J. Gazulla, M. Martinez-Tames, J. Infante and A. Jordanova (2017). “Intermediate Charcot-Marie-Tooth disease: an electrophysiological reappraisal and systematic review.” J Neurol.

Bernard-Marissal, N., G. van Hameren, M. Juneja, C. Pellegrino, L. Louhivuori, L. Bartesaghi, C. Rochat, O. El Mansour, J. J. Medard, M. Croisier, C. Maclachlan, O. Poirot, P. Uhlen, V. Timmerman, N. Tricaud, B. L. Schneider and R. Chrast (2019). “Altered interplay between endoplasmic reticulum and mitochondria in Charcot-Marie-Tooth type 2A neuropathy.” Proc Natl Acad Sci U S A 116(6): 2328–2337.

Birling, M. C., A. Yoshiki, D. J. Adams, S. Ayabe, A. L. Beaudet, J. Bottomley, A. Bradley, S. D. M. Brown, A. Burger, W. Bushell, F. Chiani, H. G. Chin, S. Christou, G. F. Codner, F. J. DeMayo, M. E. Dickinson, B. Doe, L. R. Donahue, M. D. Fray, A. Gambadoro, X. Gao, M. Gertsenstein, A. Gomez-Segura, L. O. Goodwin, J. D. Heaney, Y. Herault, M. H. de Angelis, S. T. Jiang, M. J. Justice, P. Kasparek, R. E. King, R. Kuhn, H. Lee, Y. J. Lee, Z. Liu, K. C. K. Lloyd, I. Lorenzo, A. M. Mallon, C. McKerlie, T. F. Meehan, V. M. Fuentes, S. Newman, L. M. J. Nutter, G. T. Oh, G. Pavlovic, R. Ramirez-Solis, B. Rosen, E. J. Ryder, L. A. Santos, J. Schick, J. R. Seavitt, R. Sedlacek, C. Seisenberger, J. K. Seong, W. C. Skarnes, T. Sorg, K. P. Steel, M. Tamura, G. P. Tocchini-Valentini, C. L. Wang, H. Wardle-Jones, M. Wattenhofer-Donze, S. Wells, M. V. Wiles, B. J. Willis, J. A. Wood, W. Wurst, Y. Xu, C. International Mouse Phenotyping, L. Teboul and S. A. Murray (2021). “A resource of targeted mutant mouse lines for 5,061 genes.” Nat Genet 53(4): 416–419.

Bos, R., K. Rihan, P. Quintana, L. El-Bazzal, N. Bernard-Marissal, N. Da Silva, R. Jabbour, A. Megarbane, M. Bartoli, F. Brocard and V. Delague (2022). “Altered action potential waveform and shorter axonal initial segment in hiPSC-derived motor neurons with mutations in VRK1.” Neurobiol Dis 164: 105609.

Bosco, L., Y. M. Falzone and S. C. Previtali (2021). “Animal Models as a Tool to Design Therapeutical Strategies for CMT-like Hereditary Neuropathies.” Brain Sci 11(9).

Brooks, S. P. and S. B. Dunnett (2009). “Tests to assess motor phenotype in mice: a user’s guide.” Nat Rev Neurosci 10(7): 519–529.

Canettieri, G., L. Di Marcotullio, A. Greco, S. Coni, L. Antonucci, P. Infante, L. Pietrosanti, E. De Smaele, E. Ferretti, E. Miele, M. Pelloni, G. De Simone, E. M. Pedone, P. Gallinari, A. Giorgi, C. Steinkuhler, L. Vitagliano, C. Pedone, M. E. Schinin, I. Screpanti and A. Gulino (2010). “Histone deacetylase and Cullin3-REN(KCTD11) ubiquitin ligase interplay regulates Hedgehog signalling through Gli acetylation.” Nat Cell Biol 12(2): 132–142.

Carrard, J. and F. Lejeune (2023). “Nonsense-mediated mRNA decay, a simplified view of a complex mechanism.” BMB Rep 56(12): 625–632.

Chen, X., G. Wang, Y. Zhang, M. Dayhoff-Brannigan, N. L. Diny, M. Zhao, G. He, C. N. Sing, K. A. Metz, Z. D. Stolp, A. Aouacheria, W. C. Cheng, J. M. Hardwick and X. Teng (2018). “Whi2 is a conserved negative regulator of TORC1 in response to low amino acids.” PLoS Genet 14(8): e1007592.

contributors, P. (2025). “PFMG2025-integrating genomic medicine into the national healthcare system in France.” Lancet Reg Health Eur 50: 101183.

Correale, S., L. Pirone, L. Di Marcotullio, E. De Smaele, A. Greco, D. Mazza, M. Moretti, V. Alterio, L. Vitagliano, S. Di Gaetano, A. Gulino and E. M. Pedone (2011). “Molecular organization of the cullin E3 ligase adaptor KCTD11.” Biochimie 93(4): 715–724.

Cusack, B. P., P. F. Arndt, L. Duret and H. Roest Crollius (2011). “Preventing dangerous nonsense: selection for robustness to transcriptional error in human genes.” PLoS Genet 7(10): e1002276.

Danecek, P., J. K. Bonfield, J. Liddle, J. Marshall, V. Ohan, M. O. Pollard, A. Whitwham, T. Keane, S. A. McCarthy, R. M. Davies and H. Li (2021). “Twelve years of SAMtools and BCFtools.” Gigascience 10(2).

De Grado, A., M. Serio, P. Saveri, C. Pisciotta and D. Pareyson (2025). “Charcot-Marie-Tooth disease: a review of clinical developments and its management - What’s new in 2025?” Expert Rev Neurother: 1–16.

De Smaele, E., L. Di Marcotullio, M. Moretti, M. Pelloni, M. A. Occhione, P. Infante, D. Cucchi, A. Greco, L. Pietrosanti, J. Todorovic, S. Coni, G. Canettieri, E. Ferretti, R. Bei, M. Maroder, I. Screpanti and A. Gulino (2011). “Identification and characterization of KCASH2 and KCASH3, 2 novel Cullin3 adaptors suppressing histone deacetylase and Hedgehog activity in medulloblastoma.” Neoplasia 13(4): 374–385.

Delague, V., C. Bareil, S. Tuffery, P. Bouvagnet, E. Chouery, S. Koussa, T. Maisonobe, J. Loiselet, A. Megarbane and M. Claustres (2000). “Mapping of a new locus for autosomal recessive demyelinating Charcot-Marie-Tooth disease to 19q13.1-13.3 in a large consanguineous Lebanese family: exclusion of MAG as a candidate gene.” Am J Hum Genet 67(1): 236–243.

Denis, H., R. Deplus, P. Putmans, M. Yamada, R. Metivier and F. Fuks (2009). “Functional connection between deimination and deacetylation of histones.” Mol Cell Biol 29(18): 4982–4993.

Desmet, F. O., D. Hamroun, M. Lalande, G. Collod-Beroud, M. Claustres and C. Beroud (2009). “Human Splicing Finder: an online bioinformatics tool to predict splicing signals.” Nucleic Acids Res 37(9): e67.

Desvignes, J. P., M. Bartoli, V. Delague, M. Krahn, M. Miltgen, C. Beroud and D. Salgado (2018). “VarAFT: a variant annotation and filtration system for human next generation sequencing data.” Nucleic Acids Res. 46(W1): W545–W553.

Di Marcotullio, L., E. Ferretti, E. De Smaele, B. Argenti, C. Mincione, F. Zazzeroni, R. Gallo, L. Masuelli, M. Napolitano, M. Maroder, A. Modesti, F. Giangaspero, I. Screpanti, E. Alesse and A. Gulino (2004). “REN(KCTD11) is a suppressor of Hedgehog signaling and is deleted in human medulloblastoma.” Proc Natl Acad Sci U S A 101(29): 10833–10838.

Dobin, A. and T. R. Gingeras (2015). “Mapping RNA-seq Reads with STAR.” Curr Protoc Bioinformatics 51: 11 14 11–11 14 19.

El-Bazzal, L., A. Ghata, C. Esteve, J. Gadacha, P. Quintana, C. Castro, N. Roeckel-Trevisiol, F. Lembo, N. Lenfant, A. Megarbane, J. P. Borg, N. Levy, M. Bartoli, Y. Poitelon, P. L. Roubertoux, V. Delague and N. Bernard-Marissal (2023). “Imbalance of NRG1-ERBB2/3 signalling underlies altered myelination in Charcot-Marie-Tooth disease 4H.” Brain 146(5): 1844–1858.

El-Bazzal, L., K. Rihan, N. Bernard-Marissal, C. Castro, E. Chouery-Khoury, J. P. Desvignes, A. Atkinson, K. Bertaux, S. Koussa, N. Levy, M. Bartoli, A. Megarbane, R. Jabbour and V. Delague (2019). “Loss of Cajal bodies in motor neurons from patients with novel mutations in VRK1.” Hum Mol Genet 28(14): 2378–2394.

Escamilla, C. O., I. Filonova, A. K. Walker, Z. X. Xuan, R. Holehonnur, F. Espinosa, S. Liu, S. B. Thyme, I. A. Lopez-Garcia, D. B. Mendoza, N. Usui, J. Ellegood, A. J. Eisch, G. Konopka, J. P. Lerch, A. F. Schier, H. E. Speed and C. M. Powell (2017). “Kctd13 deletion reduces synaptic transmission via increased RhoA.” Nature 551(7679): 227–231.

Farhan, S. M., L. M. Murphy, J. F. Robinson, J. Wang, V. M. Siu, C. A. Rupar, A. N. Prasad, F. C. Consortium and R. A. Hegele (2014). “Linkage analysis and exome sequencing identify a novel mutation in KCTD7 in patients with progressive myoclonus epilepsy with ataxia.” Epilepsia 55(9): e106–111.

Fattahi, Z., M. Beheshtian, M. Mohseni, H. Poustchi, E. Sellars, S. H. Nezhadi, A. Amini, S. Arzhangi, K. Jalalvand, P. Jamali, Z. Mohammadi, B. Davarnia, P. Nikuei, M. Oladnabi, A. Mohammadzadeh, E. Zohrehvand, A. Nejatizadeh, M. Shekari, M. Bagherzadeh, E. Shamsi-Gooshki, S. Borno, B. Timmermann, A. Haghdoost, R. Najafipour, H. R. Khorram Khorshid, K. Kahrizi, R. Malekzadeh, M. R. Akbari and H. Najmabadi (2019). “Iranome: A catalog of genomic variations in the Iranian population.” Hum Mutat 40(11): 1968–1984.

Figlia, G., D. Gerber and U. Suter (2018). “Myelination and mTOR.” Glia 66(4): 693–707.

Gallo, R., F. Zazzeroni, E. Alesse, C. Mincione, U. Borello, P. Buanne, R. D’Eugenio, A. R. Mackay, B. Argenti, R. Gradini, M. A. Russo, M. Maroder, G. Cossu, L. Frati, I. Screpanti and A. Gulino (2002). “REN: a novel, developmentally regulated gene that promotes neural cell differentiation.” J Cell Biol 158(4): 731–740.

Goddard, T. D., C. C. Huang, E. C. Meng, E. F. Pettersen, G. S. Couch, J. H. Morris and T. E. Ferrin (2018). “UCSF ChimeraX: Meeting modern challenges in visualization and analysis.” Protein Sci 27(1): 14–25.

Golzio, C., J. Willer, M. E. Talkowski, E. C. Oh, Y. Taniguchi, S. Jacquemont, A. Reymond, M. Sun, A. Sawa, J. F. Gusella, A. Kamiya, J. S. Beckmann and N. Katsanis (2012). “KCTD13 is a major driver of mirrored neuroanatomical phenotypes of the 16p11.2 copy number variant.” Nature 485(7398): 363–367.

Gomez-Sanchez, J. A., L. Carty, M. Iruarrizaga-Lejarreta, M. Palomo-Irigoyen, M. Varela-Rey, M. Griffith, J. Hantke, N. Macias-Camara, M. Azkargorta, I. Aurrekoetxea, V. G. De Juan, H. B. Jefferies, P. Aspichueta, F. Elortza, A. M. Aransay, M. L. Martinez-Chantar, F. Baas, J. M. Mato, R. Mirsky, A. Woodhoo and K. R. Jessen (2015). “Schwann cell autophagy, myelinophagy, initiates myelin clearance from injured nerves.” J Cell Biol 210(1): 153–168.

Groza, T., F. L. Gomez, H. H. Mashhadi, V. Munoz-Fuentes, O. Gunes, R. Wilson, P. Cacheiro, A. Frost, P. Keskivali-Bond, B. Vardal, A. McCoy, T. K. Cheng, L. Santos, S. Wells, D. Smedley, A. M. Mallon and H. Parkinson (2023). “The International Mouse Phenotyping Consortium: comprehensive knockout phenotyping underpinning the study of human disease.” Nucleic Acids Res 51(D1): D1038–D1045.

Harding, A. E. and P. K. Thomas (1980). “The clinical features of hereditary motor and sensory neuropathy types I and II.” Brain 103(2): 259–280.

Higuchi, Y. and H. Takashima (2023). “Clinical genetics of Charcot-Marie-Tooth disease.” J Hum Genet 68(3): 199–214.

Jacob, C., C. N. Christen, J. A. Pereira, C. Somandin, A. Baggiolini, P. Lotscher, M. Ozcelik, N. Tricaud, D. Meijer, T. Yamaguchi, P. Matthias and U. Suter (2011). “HDAC1 and HDAC2 control the transcriptional program of myelination and the survival of Schwann cells.” Nat Neurosci 14(4): 429–436.

Jacob, C., P. Lotscher, S. Engler, A. Baggiolini, S. Varum Tavares, V. Brugger, N. John, S. Buchmann-Moller, P. L. Snider, S. J. Conway, T. Yamaguchi, P. Matthias, L. Sommer, N. Mantei and U. Suter (2014). “HDAC1 and HDAC2 control the specification of neural crest cells into peripheral glia.” J Neurosci 34(17): 6112–6122.

Jaganathan, K., S. Kyriazopoulou Panagiotopoulou, J. F. McRae, S. F. Darbandi, D. Knowles, Y. I. Li, J. A. Kosmicki, J. Arbelaez, W. Cui, G. B. Schwartz, E. D. Chow, E. Kanterakis, H. Gao, A. Kia, S. Batzoglou, S. J. Sanders and K. K. Farh (2019). “Predicting Splicing from Primary Sequence with Deep Learning.” Cell 176(3): 535–548 e524.

Jobling, R. K., M. Assoum, O. Gakh, S. Blaser, J. A. Raiman, C. Mignot, E. Roze, A. Durr, A. Brice, N. Levy, C. Prasad, T. Paton, A. D. Paterson, N. M. Roslin, C. R. Marshall, J. P. Desvignes, N. Roeckel-Trevisiol, S. W. Scherer, G. A. Rouleau, A. Megarbane, G. Isaya, V. Delague and G. Yoon (2015). “PMPCA mutations cause abnormal mitochondrial protein processing in patients with non-progressive cerebellar ataxia.” Brain 138(Pt 6): 1505–1517.

Jumper, J., R. Evans, A. Pritzel, T. Green, M. Figurnov, O. Ronneberger, K. Tunyasuvunakool, R. Bates, A. Zidek, A. Potapenko, A. Bridgland, C. Meyer, S. A. A. Kohl, A. J. Ballard, A. Cowie, B. Romera-Paredes, S. Nikolov, R. Jain, J. Adler, T. Back, S. Petersen, D. Reiman, E. Clancy, M. Zielinski, M. Steinegger, M. Pacholska, T. Berghammer, S. Bodenstein, D. Silver, O. Vinyals, A. W. Senior, K. Kavukcuoglu, P. Kohli and D. Hassabis (2021). “Highly accurate protein structure prediction with AlphaFold.” Nature 596(7873): 583–589.

Kaiser, T., H. M. Allen, O. Kwon, B. Barak, J. Wang, Z. He, M. Jiang and G. Feng (2021). “MyelTracer: A Semi-Automated Software for Myelin g-Ratio Quantification.” eNeuro 8(4).

Karousis, E. D. and O. Muhlemann (2019). “Nonsense-Mediated mRNA Decay Begins Where Translation Ends.” Cold Spring Harb Perspect Biol 11(2).

Kircher, M., D. M. Witten, P. Jain, B. J. O’Roak, G. M. Cooper and J. Shendure (2014). “A general framework for estimating the relative pathogenicity of human genetic variants.” Nature Genetics 46(3): 310–315.

Kousi, M., V. Anttila, A. Schulz, S. Calafato, E. Jakkula, E. Riesch, L. Myllykangas, H. Kalimo, M. Topcu, S. Gokben, F. Alehan, J. R. Lemke, M. Alber, A. Palotie, O. Kopra and A. E. Lehesjoki (2012). “Novel mutations consolidate KCTD7 as a progressive myoclonus epilepsy gene.” J Med Genet 49(6): 391–399.

Krabichler, B., K. Rostasy, M. Baumann, D. Karall, S. Scholl-Burgi, C. Schwarzer, K. Gautsch, A. Spreiz, D. Kotzot, J. Zschocke, C. Fauth and E. Haberlandt (2012). “Novel mutation in potassium channel related gene KCTD7 and progressive myoclonic epilepsy.” Ann Hum Genet 76(4): 326–331.

Kumar, P., S. Henikoff and P. C. Ng (2009). “Predicting the effects of coding non-synonymous variants on protein function using the SIFT algorithm.” Nature Protocols 4(7): 1073–1081.

Lander, E. S. and D. Botstein (1987). “Homozygosity mapping: a way to map human recessive traits with the DNA of inbred children.” Science 236(4808): 1567–1570.

Laura, M., M. Pipis, A. M. Rossor and M. M. Reilly (2019). “Charcot-Marie-Tooth disease and related disorders: an evolving landscape.” Curr Opin Neurol 32(5): 641–650.

Li, H. (2013). “Aligning sequence reads, clone sequences and assembly contigs with BWA-MEM.” arXiv.

Li, H. and R. Durbin (2009). “Fast and accurate short read alignment with Burrows-Wheeler transform.” Bioinformatics (Oxford, England) 25(14): 1754–1760.

Lorzadeh, S., L. Kohan, S. Ghavami and N. Azarpira (2021). “Autophagy and the Wnt signaling pathway: A focus on Wnt/beta-catenin signaling.” Biochim Biophys Acta Mol Cell Res 1868(3): 118926.

Love, M. I., W. Huber and S. Anders (2014). “Moderated estimation of fold change and dispersion for RNA-seq data with DESeq2.” Genome Biol 15(12): 550.

Luningschror, P., B. Binotti, B. Dombert, P. Heimann, A. Perez-Lara, C. Slotta, N. Thau-Habermann, R. v. C. C, F. Karl, M. Damme, A. Horowitz, I. Maystadt, A. Fuchtbauer, E. M. Fuchtbauer, S. Jablonka, R. Blum, N. Uceyler, S. Petri, B. Kaltschmidt, R. Jahn, C. Kaltschmidt and M. Sendtner (2017). “Plekhg5-regulated autophagy of synaptic vesicles reveals a pathogenic mechanism in motoneuron disease.” Nat Commun 8(1): 678.

Luningschror, P., C. Slotta, P. Heimann, M. Briese, U. M. Weikert, B. Massih, S. Appenzeller, M. Sendtner, C. Kaltschmidt and B. Kaltschmidt (2020). “Absence of Plekhg5 Results in Myelin Infoldings Corresponding to an Impaired Schwann Cell Autophagy, and a Reduced T-Cell Infiltration Into Peripheral Nerves.” Front Cell Neurosci 14: 185.

Makoukji, J., G. Shackleford, D. Meffre, J. Grenier, P. Liere, J. M. Lobaccaro, M. Schumacher and C. Massaad (2011). “Interplay between LXR and Wnt/beta-catenin signaling in the negative regulation of peripheral myelin genes by oxysterols.” J Neurosci 31(26): 9620–9629.

Maurice, M. M. and S. Angers (2025). “Mechanistic insights into Wnt-beta-catenin pathway activation and signal transduction.” Nat Rev Mol Cell Biol 26(5): 371–388.

Maystadt, I., R. Rezsohazy, M. Barkats, S. Duque, P. Vannuffel, S. Remacle, B. Lambert, M. Najimi, E. Sokal, A. Munnich, L. Viollet and C. Verellen-Dumoulin (2007). “The nuclear factor kappaB-activator gene PLEKHG5 is mutated in a form of autosomal recessive lower motor neuron disease with childhood onset.” Am J Hum Genet 81(1): 67–76.

McKenna, A., M. Hanna, E. Banks, A. Sivachenko, K. Cibulskis, A. Kernytsky, K. Garimella, D. Altshuler, S. Gabriel, M. Daly and M. A. DePristo (2010). “The Genome Analysis Toolkit: a MapReduce framework for analyzing next-generation DNA sequencing data.” Genome Research 20(9): 1297–1303.

Mencacci, N. E., I. Rubio-Agusti, A. Zdebik, F. Asmus, M. H. Ludtmann, M. Ryten, V. Plagnol, A. K. Hauser, S. Bandres-Ciga, C. Bettencourt, P. Forabosco, D. Hughes, M. M. Soutar, K. Peall, H. R. Morris, D. Trabzuni, M. Tekman, H. C. Stanescu, R. Kleta, M. Carecchio, G. Zorzi, N. Nardocci, B. Garavaglia, E. Lohmann, A. Weissbach, C. Klein, J. Hardy, A. M. Pittman, T. Foltynie, A. Y. Abramov, T. Gasser, K. P. Bhatia and N. W. Wood (2015). “A missense mutation in KCTD17 causes autosomal dominant myoclonus-dystonia.” Am J Hum Genet 96(6): 938–947.

Metz, K. A., X. Teng, I. Coppens, H. M. Lamb, B. E. Wagner, J. A. Rosenfeld, X. Chen, Y. Zhang, H. J. Kim, M. E. Meadow, T. S. Wang, E. D. Haberlandt, G. W. Anderson, E. Leshinsky-Silver, W. Bi, T. C. Markello, M. Pratt, N. Makhseed, A. Garnica, N. R. Danylchuk, T. A. Burrow, P. Jayakar, D. McKnight, S. Agadi, H. Gbedawo, C. Stanley, M. Alber, I. Prehl, K. Peariso, M. T. Ong, S. R. Mordekar, M. J. Parker, D. Crooks, P. B. Agrawal, G. T. Berry, T. Loddenkemper, Y. Yang, G. H. B. Maegawa, A. Aouacheria, J. G. Markle, J. A. Wohlschlegel, A. L. Hartman and J. M. Hardwick (2018). “KCTD7 deficiency defines a distinct neurodegenerative disorder with a conserved autophagy-lysosome defect.” Ann Neurol 84(5): 766–780.

Nicholson, G. and S. Myers (2006). “Intermediate forms of Charcot-Marie-Tooth neuropathy: a review.” Neuromolecular Med 8(1-2): 123–130.

Pareyson, D. and C. Marchesi (2009). “Diagnosis, natural history, and management of Charcot-Marie-Tooth disease.” Lancet Neurol 8(7): 654–667.

Pertea, M., G. M. Pertea, C. M. Antonescu, T. C. Chang, J. T. Mendell and S. L. Salzberg (2015). “StringTie enables improved reconstruction of a transcriptome from RNA-seq reads.” Nat Biotechnol 33(3): 290–295.

Petherick, K. J., A. C. Williams, J. D. Lane, P. Ordonez-Moran, J. Huelsken, T. J. Collard, H. J. Smartt, J. Batson, K. Malik, C. Paraskeva and A. Greenhough (2013). “Autolysosomal beta-catenin degradation regulates Wnt-autophagy-p62 crosstalk.” EMBO J 32(13): 1903–1916.

Pipis, M., A. M. Rossor, M. Laura and M. M. Reilly (2019). “Next-generation sequencing in Charcot-Marie-Tooth disease: opportunities and challenges.” Nat Rev Neurol 15(11): 644–656.

Pisciotta, C. and M. E. Shy (2018). “Neuropathy.” Handb Clin Neurol 148: 653–665.

Pisciotta, C. and M. E. Shy (2023). “Hereditary neuropathy.” Handb Clin Neurol 195: 609–617.

Poitelon, Y. and M. L. Feltri (2018). “The Pseudopod System for Axon-Glia Interactions: Stimulation and Isolation of Schwann Cell Protrusions that Form in Response to Axonal Membranes.” Methods Mol Biol 1739: 233–253.

Salgado, D., J.-P. Desvignes, G. Rai, A. Blanchard, M. Miltgen, A. Pinard, N. Lévy, G. Collod-Béroud and C. Béroud (2016). “UMD-Predictor: A High-Throughput Sequencing Compliant System for Pathogenicity Prediction of any Human cDNA Substitution.” Human Mutation 37(5): 439–446.

Schulz, A., C. Walther, H. Morrison and R. Bauer (2014). “In vivo electrophysiological measurements on mouse sciatic nerves.” J Vis Exp(86).

Schwarz, J. M., C. Rödelsperger, M. Schuelke and D. Seelow (2010). “MutationTaster evaluates disease-causing potential of sequence alterations.” Nature Methods 7(8): 575–576.

Schwenk, J., M. Metz, G. Zolles, R. Turecek, T. Fritzius, W. Bildl, E. Tarusawa, A. Kulik, A. Unger, K. Ivankova, R. Seddik, J. Y. Tiao, M. Rajalu, J. Trojanova, V. Rohde, M. Gassmann, U. Schulte, B. Fakler and B. Bettler (2010). “Native GABA(B) receptors are heteromultimers with a family of auxiliary subunits.” Nature 465(7295): 231–235.

Scott, E. M., A. Halees, Y. Itan, E. G. Spencer, Y. He, M. A. Azab, S. B. Gabriel, A. Belkadi, B. Boisson, L. Abel, A. G. Clark, C. Greater Middle East Variome, F. S. Alkuraya, J. L. Casanova and J. G. Gleeson (2016). “Characterization of Greater Middle Eastern genetic variation for enhanced disease gene discovery.” Nat Genet 48(9): 1071–1076.

Skre, H. (1974). “Genetic and clinical aspects of Charcot-Marie-Tooth’s disease.” Clin Genet 6(2): 98–118.

Tawk, M., J. Makoukji, M. Belle, C. Fonte, A. Trousson, T. Hawkins, H. Li, S. Ghandour, M. Schumacher and C. Massaad (2011). “Wnt/beta-catenin signaling is an essential and direct driver of myelin gene expression and myelinogenesis.” J Neurosci 31(10): 3729–3742.

Tong, R., B. Yang, H. Xiao, C. Peng, W. Hu, X. Weng, S. Cheng, C. Du, Z. Lv, C. Ding, L. Zhou, H. Xie, J. Wu and S. Zheng (2017). “KCTD11 inhibits growth and metastasis of hepatocellular carcinoma through activating Hippo signaling.” Oncotarget 8(23): 37717–37729.

Wang, K., M. Li and H. Hakonarson (2010). “ANNOVAR: functional annotation of genetic variants from high-throughput sequencing data.” Nucleic Acids Research 38(16): e164.

Wang, Y., D. Argiles-Castillo, E. I. Kane, A. Zhou and D. E. Spratt (2020). “HECT E3 ubiquitin ligases - emerging insights into their biological roles and disease relevance.” J Cell Sci 133(7).

Willer, C. J., E. K. Speliotes, R. J. Loos, S. Li, C. M. Lindgren, I. M. Heid, S. I. Berndt, A. L. Elliott, A. U. Jackson, C. Lamina, G. Lettre, N. Lim, H. N. Lyon, S. A. McCarroll, K. Papadakis, L. Qi, J. C. Randall, R. M. Roccasecca, S. Sanna, P. Scheet, M. N. Weedon, E. Wheeler, J. H. Zhao, L. C. Jacobs, I. Prokopenko, N. Soranzo, T. Tanaka, N. J. Timpson, P. Almgren, A. Bennett, R. N. Bergman, S. A. Bingham, L. L. Bonnycastle, M. Brown, N. P. Burtt, P. Chines, L. Coin, F. S. Collins, J. M. Connell, C. Cooper, G. D. Smith, E. M. Dennison, P. Deodhar, P. Elliott, M. R. Erdos, K. Estrada, D. M. Evans, L. Gianniny, C. Gieger, C. J. Gillson, C. Guiducci, R. Hackett, D. Hadley, A. S. Hall, A. S. Havulinna, J. Hebebrand, A. Hofman, B. Isomaa, K. B. Jacobs, T. Johnson, P. Jousilahti, Z. Jovanovic, K. T. Khaw, P. Kraft, M. Kuokkanen, J. Kuusisto, J. Laitinen, E. G. Lakatta, J. Luan, R. N. Luben, M. Mangino, W. L. McArdle, T. Meitinger, A. Mulas, P. B. Munroe, N. Narisu, A. R. Ness, K. Northstone, S. O’Rahilly, C. Purmann, M. G. Rees, M. Ridderstrale, S. M. Ring, F. Rivadeneira, A. Ruokonen, M. S. Sandhu, J. Saramies, L. J. Scott, A. Scuteri, K. Silander, M. A. Sims, K. Song, J. Stephens, S. Stevens, H. M. Stringham, Y. C. Tung, T. T. Valle, C. M. Van Duijn, K. S. Vimaleswaran, P. Vollenweider, G. Waeber, C. Wallace, R. M. Watanabe, D. M. Waterworth, N. Watkins, C. Wellcome Trust Case Control, J. C. Witteman, E. Zeggini, G. Zhai, M. C. Zillikens, D. Altshuler, M. J. Caulfield, S. J. Chanock, I. S. Farooqi, L. Ferrucci, J. M. Guralnik, A. T. Hattersley, F. B. Hu, M. R. Jarvelin, M. Laakso, V. Mooser, K. K. Ong, W. H. Ouwehand, V. Salomaa, N. J. Samani, T. D. Spector, T. Tuomi, J. Tuomilehto, M. Uda, A. G. Uitterlinden, N. J. Wareham, P. Deloukas, T. M. Frayling, L. C. Groop, R. B. Hayes, D. J. Hunter, K. L. Mohlke, L. Peltonen, D. Schlessinger, D. P. Strachan, H. E. Wichmann, M. I. McCarthy, M. Boehnke, I. Barroso, G. R. Abecasis, J. N. Hirschhorn and A. T. C. Genetic Investigation of (2009). “Six new loci associated with body mass index highlight a neuronal influence on body weight regulation.” Nat Genet 41(1): 25–34.

Yang, M., Y. M. Han, Q. Han, X. Z. Rong, X. F. Liu and X. Y. Ln (2021). “KCTD11 inhibits progression of lung cancer by binding to beta-catenin to regulate the activity of the Wnt and Hippo pathways.” J Cell Mol Med 25(19): 9411–9426.

Zazzeroni, F., D. Nicosia, A. Tessitore, R. Gallo, D. Verzella, M. Fischietti, D. Vecchiotti, L. Ventura, D. Capece, A. Gulino and E. Alesse (2014). “KCTD11 tumor suppressor gene expression is reduced in prostate adenocarcinoma.” Biomed Res Int 2014: 380398.

