## Supplementary figures and images for "Bi-allelic mutations in *KCTD11* cause a new form of autosomal recessive intermediate Charcot-Marie-Tooth disease"

### Supplementary Figures 1 to 7

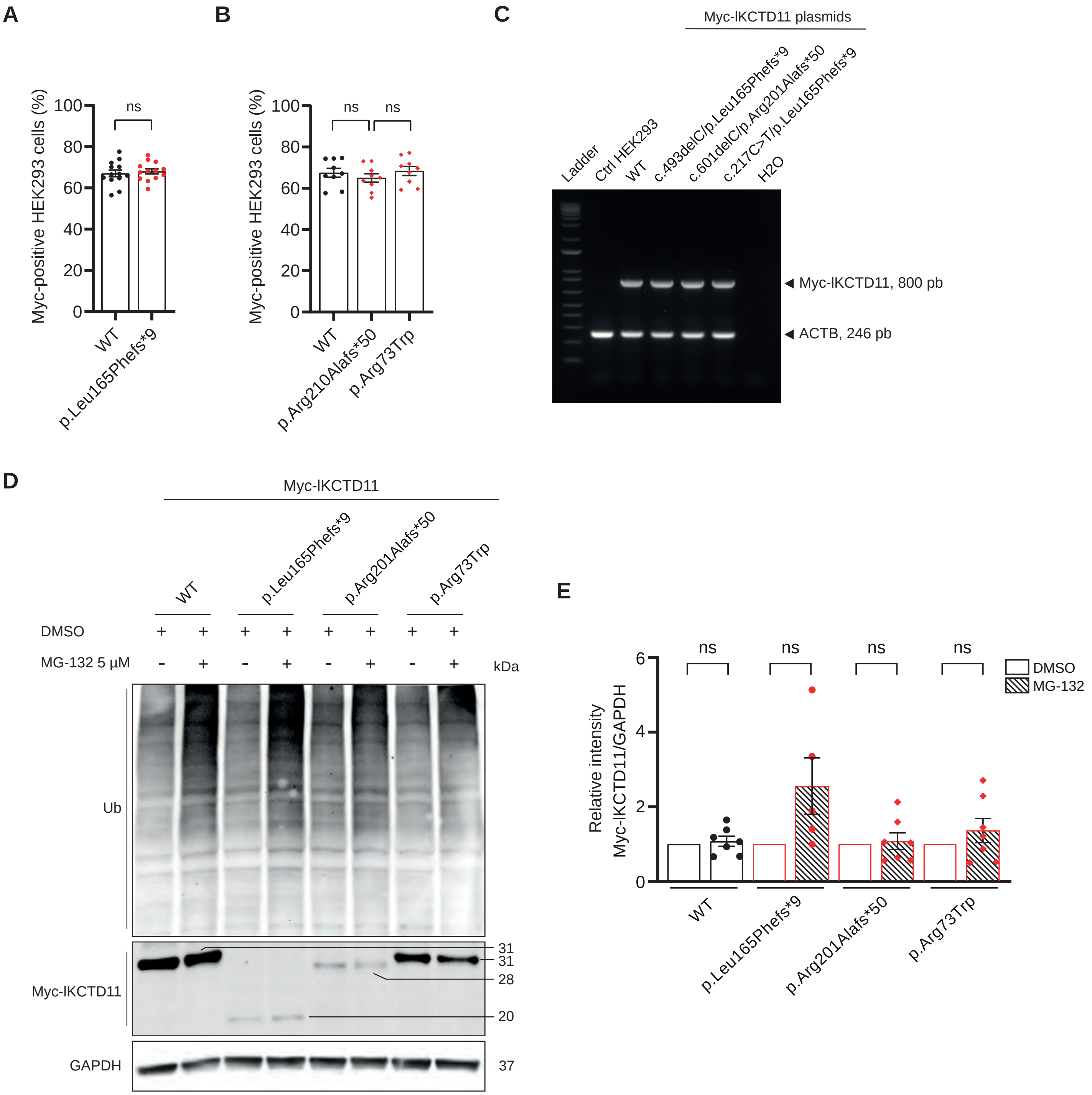

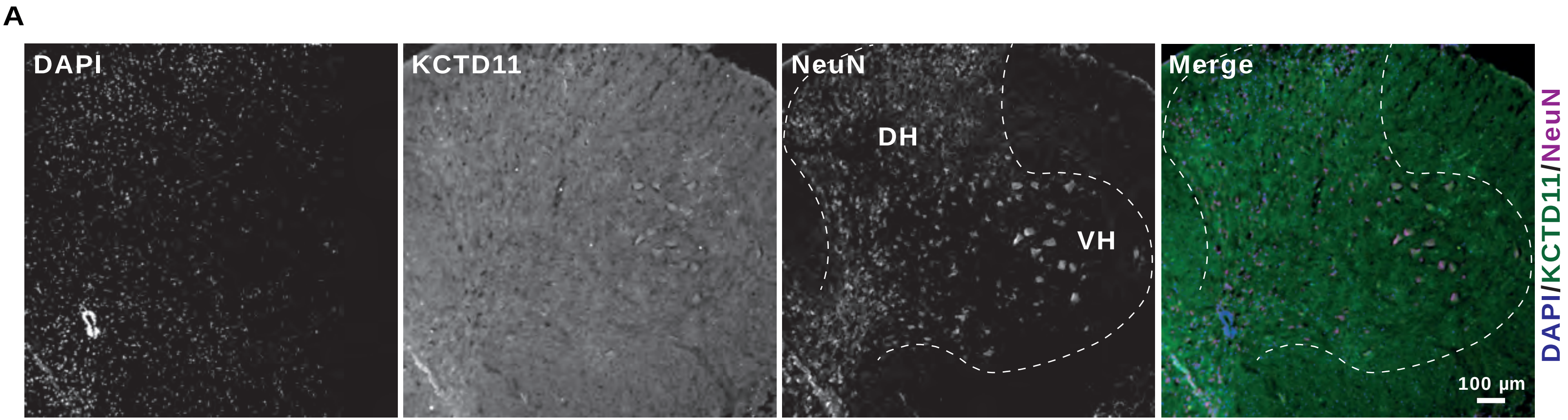

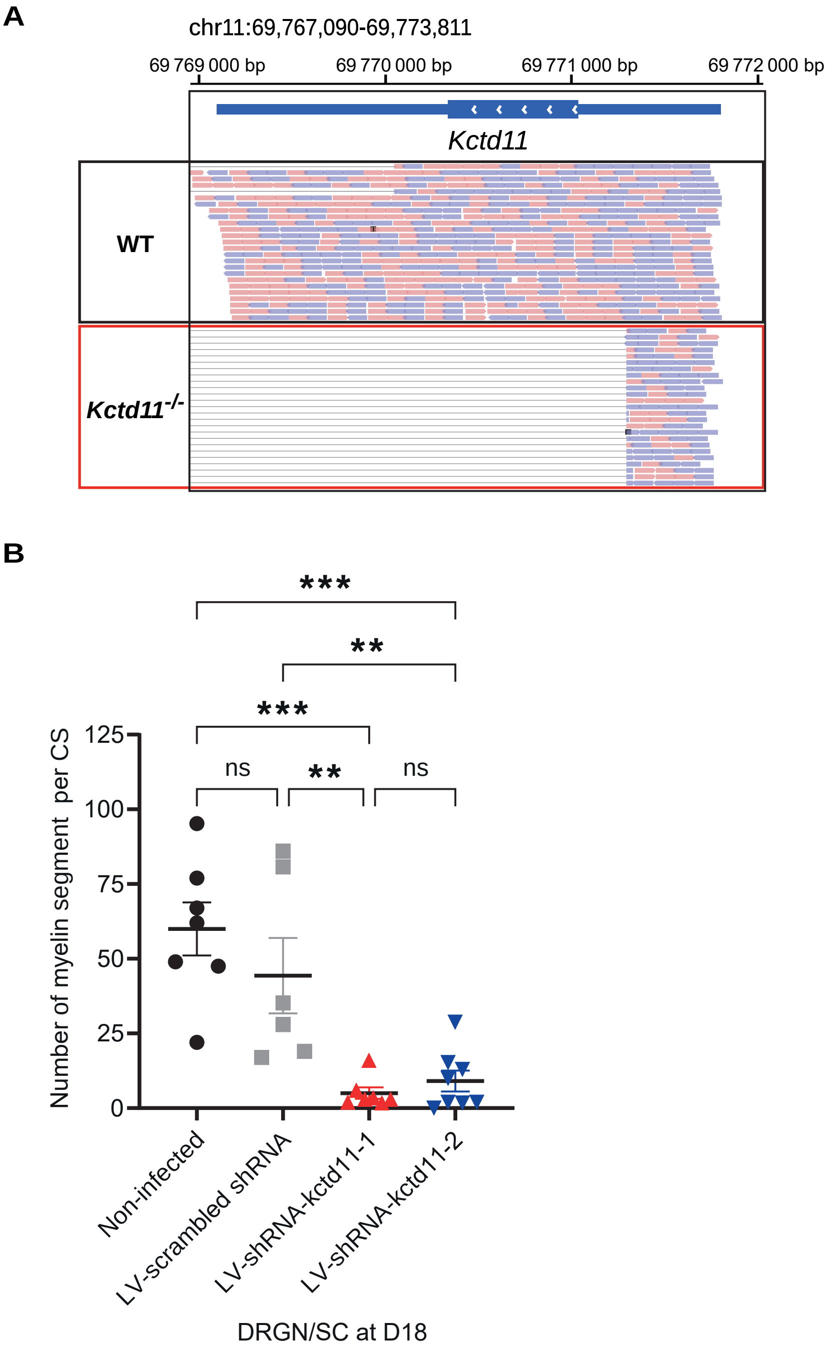

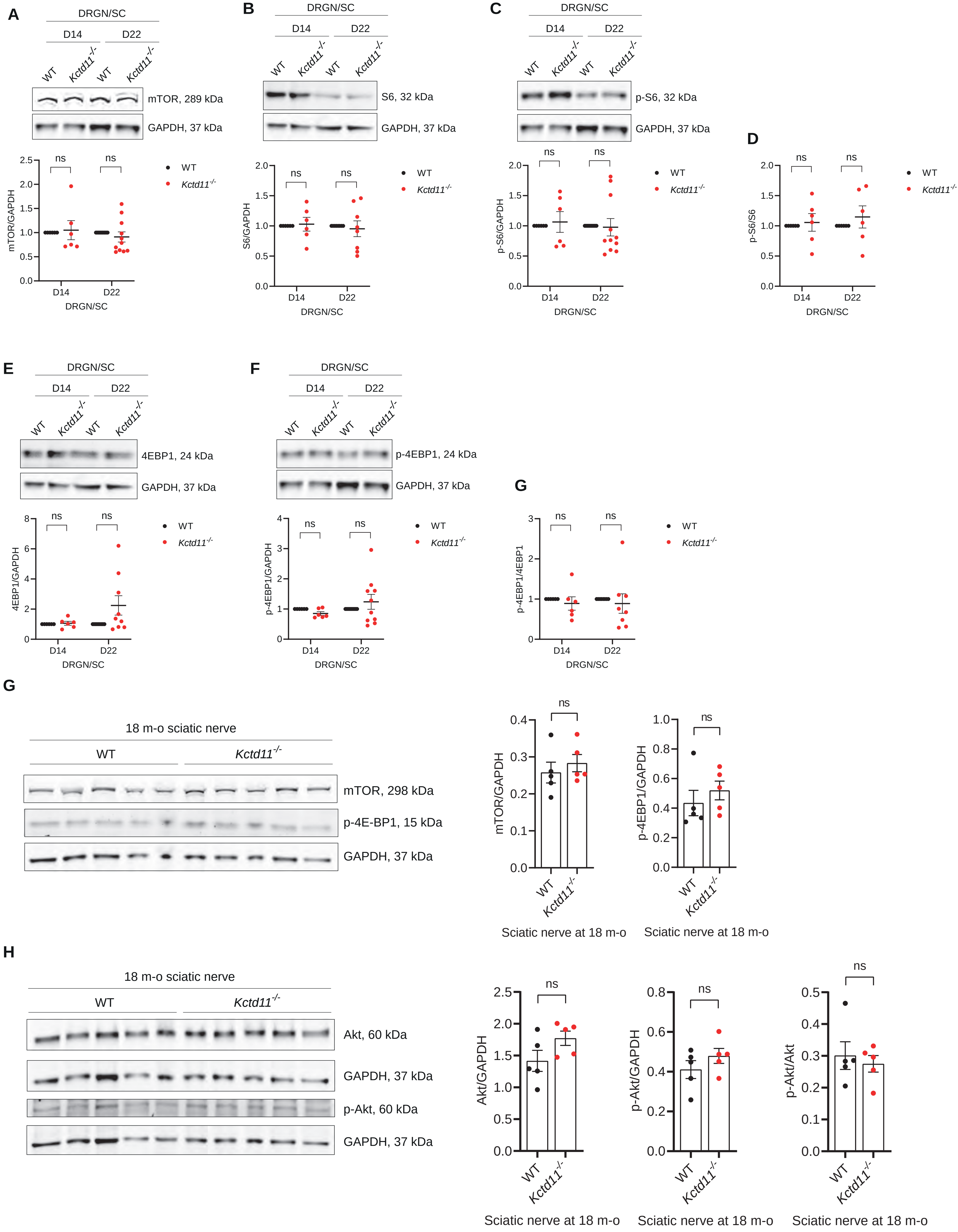

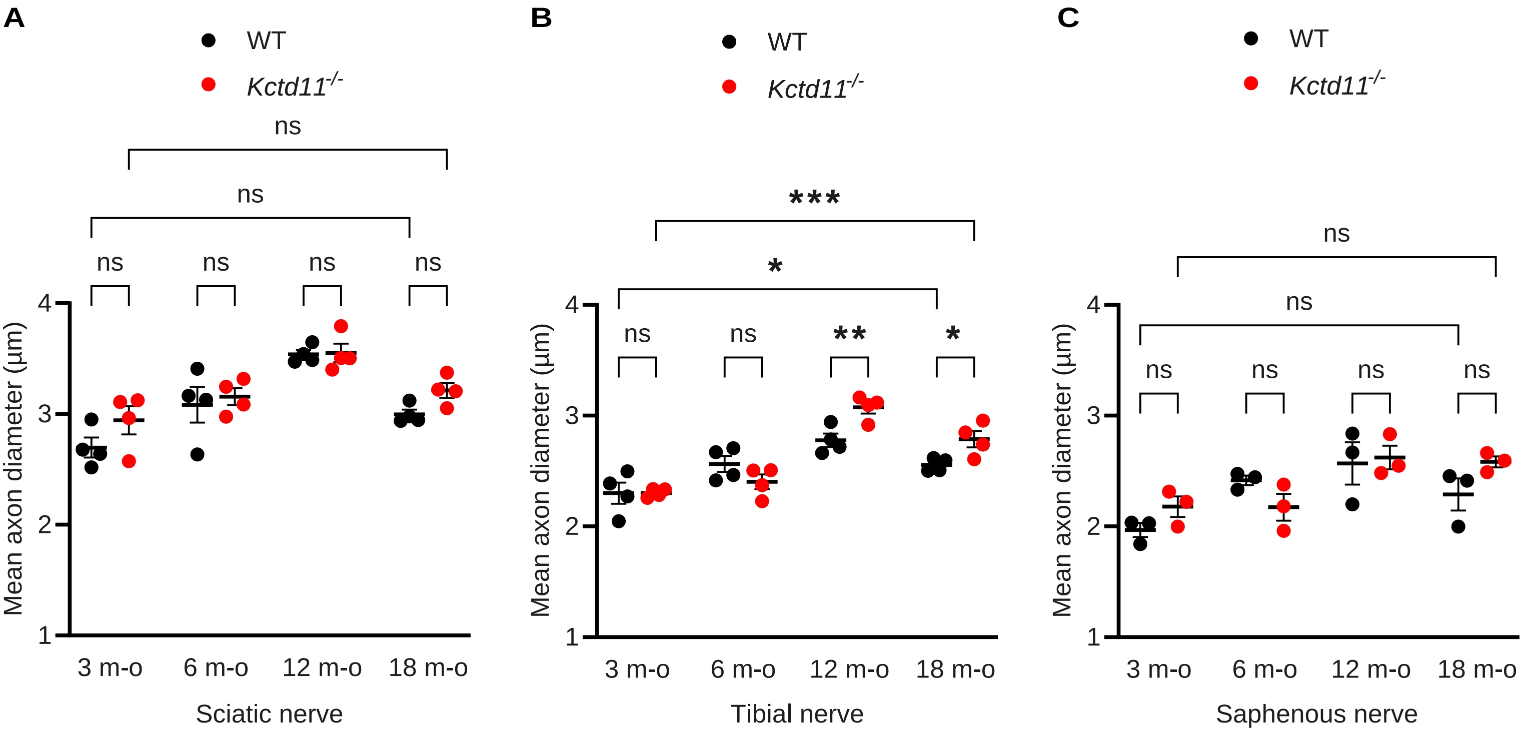

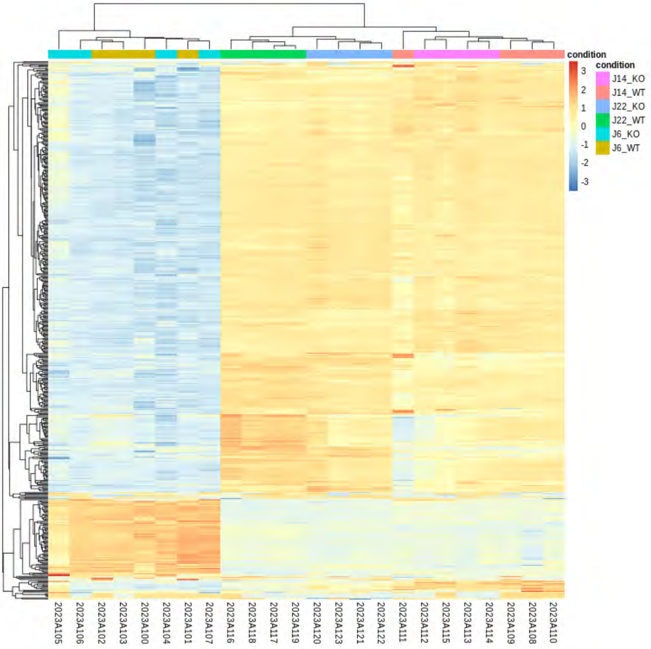

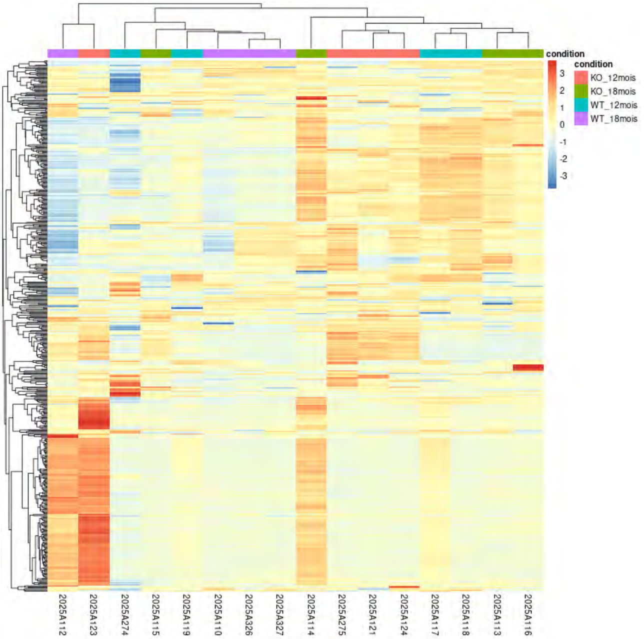

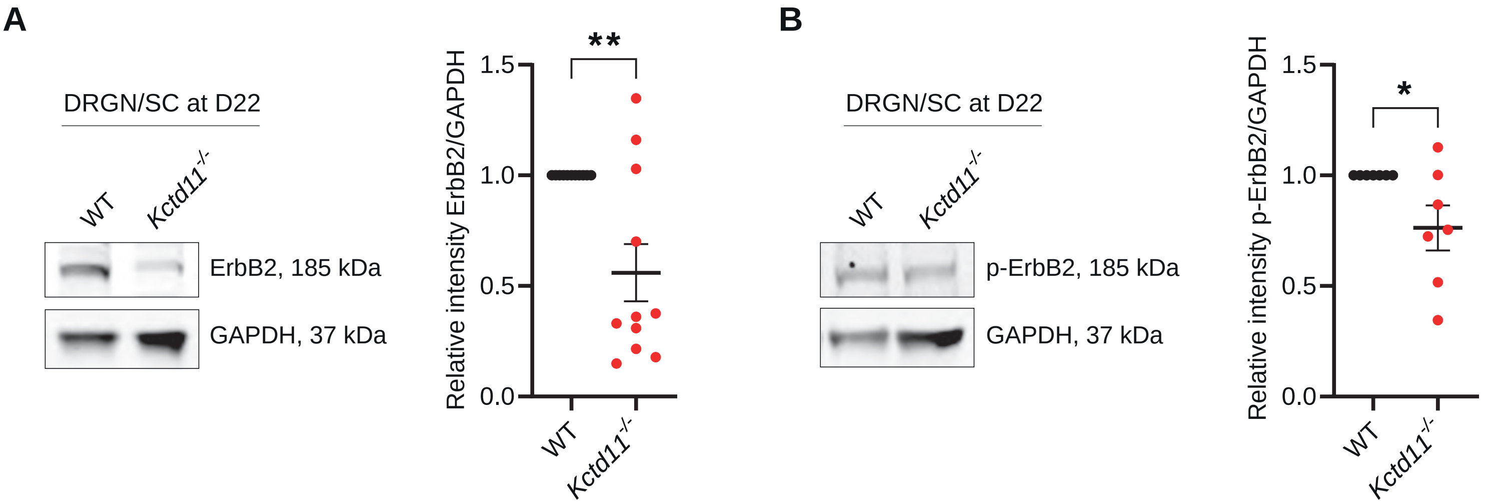
